# Chitosan induces salicylic acid local and systemically in banana plants and reduces colonization by the pathogen *Fusarium oxysporum* f. sp. cubense TR4

**DOI:** 10.1101/2024.02.27.582266

**Authors:** Federico Lopez-Moya, Jassmin Zorrilla-Fontanesi, Ana Lozano-Soria, Naia Fernandez de Larrinoa Ganado, Celia Mei Moreno González, Arnau Hernández, Ariadna Torres, Daniel Gonzalez-Silvera, Benet Gunsé, Jose Angel Lopez Jimenez, Luis V. Lopez-Llorca

**Affiliations:** Department of Marine Sciences and Applied Biology, Laboratory of Plant Pathology, University of Alicante, 03080 Alicante, Spain; Laboratory of Tropical Crop Improvement, Division of Crop Biotechnics, KU Leuven, Leuven, Belgium; Plant Physiology Laboratory, Faculty of Biosciences, Universidad Autónoma de Barcelona, Bellaterra, Spain; Department of Physiology, Faculty of Biology, Murcia, 30100 Murcia University, Spain

**Keywords:** Chitosan, root exudates, membrane depolarization, membrane fluidity, reactive oxygen species (ROS), salicylic acid, methyl salicylate, volatile organic compounds (VOCs), systemic acquired resistance (SAR), *Pochonia chlamydosporia*.

## Abstract

Banana (*Musa* spp.) is essential for food security. For its clonality is highly susceptible to pests and diseases. The wilt fungus *Fusarium oxysporum* f. sp. *cubense* Tropical Race 4 (FocTR4) threatens banana production worldwide. We notice that chitosan protects banana from biotic stress. Chitosan, a plant defence elicitor with antimicrobial activity, triggers salicylic acid (SA) and methyl salicylate biosynthesis and induces systemic acquired resistance (SAR) genes, mainly flavin-dependent monooxygenase 1 (FMO1), in banana. Chitosan promotes root exudation of SA and emission of methyl salicylate (MeSA). Banana germplasm, *Petit Naine*, *Gros Michel*, *Enzirabahima*, *Yangambi Km5* and *Foconah* banana differ in their response to chitosan. Chitosan induces, in Foc resistant *Yangambi Km5*, ca. 7-fold FMO1. Preventive chitosan treatments together with the endophytic biocontrol fungus *Pochonia chlamydosporia* strain 123 reduce (ca. 4-fold) colonization of banana roots by FocTR4. Therefore, chitosan and beneficial endophytes can help to manage sustainably FocTR4 in banana.

## Introduction

Bananas are an essential crop for food security worldwide. They are grown in most tropical and subtropical countries^1,2^. Current banana plant cultivars derive from hybrids of diploid wild species *Musa acuminata* (Colla, 1820; Genome A) and *Musa balbisiana* (Colla, 1820; Genome B)^3^. These plants are seedless triploids (AAA, AAB or ABB)^4^ and there are propagated clonally. Cavendish (AAA) is the most internationally traded banana. Its low genetic diversity makes it highly susceptible to pests and diseases. Insects (*Cosmopolites sordidus*, Germar 1824; banana weevil, BW), plant-parasitic nematodes (*Pratylenchus goodeyi*, *Helicotylenchus multicinctus*, *Radopholus similis* and *Meloidogyne* spp.), wilt fungi (*Fusarium oxysporum* f. sp. *cubense,* (Smith) Snyder and Hansen, Foc) and bacteria (*Xanthomonas campestris* pv. *musacearum*)^5,6,7^ are major biotic stresses to banana. Management of these stresses in banana is based on the use of chemicals with a high impact on the agroecosystems and environment. Current regulations restrict or ban toxic agrochemicals in banana plantations^8^. Therefore, new sustainable strategies based on biomolecules and biocontrol agents to manage agronomical pests and diseases are required.

Chitin is a key structural component of the exoskeleton of invertebrates and fungal cell walls. It is an important molecular pattern of the innate immune systems for pathogen recognition in plants, both monocots and dicots, and animals^9^. Chitosan is a linear biopolymer based on β-(1-4)-linked D-glucosamine and N-acetyl-β-D-glucosamine randomly distributed, obtained by partial deacetylation of chitin^10^. Chitosan is non-toxic to plants, animals, and mammalian cells at moderate concentrations^11,12^. It is also biodegradable and environmentally safe^13^. Chitosan kills sensitive fungi and bacteria, including important plant pathogens like *Fusarium oxysporum* f.sp radicis-lycopersici or *Xanthomonas* spp^14,15,16^. Therefore, chitosan reduces plant diseases caused by plant pathogenic microorganisms^17,18,19,20^. Chitosan antimicrobial action is mediated by plasma membrane permeabilization of microorganisms with enriched unsaturated free fatty acids plasma membranes and also related to an oxidative burst which contribute to unsaturated free fatty acids oxidation^21^.

Chitosan has also been described as an elicitor of plant defences^21,22,23,24^. For instance, chitosan induces biosynthesis of jasmonic and salicylic acid^12,25,26^. Salicylic acid (SA) methylation (MeSA) increases SA membrane permeability^27^. MeSA can be released to be used as a cue when needed^28,29^. MeSA accumulates upon stress in infected tissues and activates and amplifies systemic acquired resistance (SAR) in healthy distal plant tissues, and in surrounding plants too ^27,30,31,32,33,34^. For SAR MeSA it is converted back to SA^35^.

Chitosan also modifies root cell membranes architecture and root differentiation^12,36^. Plasma membrane of plant cells regulates nutrient exchanges and controls signal perception, inducing adaptive responses to environmental changes^37^. Plant membrane fatty acids and volatile lipoxygenase (LOX) pathway products are released after small mechanical damage^38,39,40^. Jasmonic acid, a lipid derivative, and methyl jasmonate, also affect the expression of defence related genes and the production of defence metabolites^41^. PAMP immunity involves alterations in plant membrane lipid composition^42^. This modifies membrane fluidity and generates oxylipins as a signalling compounds^43,44^. Lipids can then act as biomarkers for environmental stress^45^. Very-long-chain fatty acids (VLCFAs) are important for plant defence, since they are required for the biosynthesis of plant cuticle and sphingolipids, involved in cell death regulation^46^.

In this work we analyse the effect of chitosan applied to the rhizosphere of banana plants. Our main objective is to assess the effect of chitosan on the induction of plant defence. We characterise the effect of chitosan on the banana plant cell plasma membrane potential and fatty acid composition. We also test the effect of chitosan on banana rhizodeposition (SA and other hormones) and leaf volatilome. In this study we have used the most commercial banana cultivar *Musa acuminata* (*Petite Naine*, Cavendish AAA) besides other banana germplasm (*Yangambi km5* ITC1123, Ibota Bota AAA; *Gros Michel* ITC1122, *Gros Michel* AAA; *Enzirabahima* ITC1354, EAHB, Mutika/Lujugira AAA; *Foconah* ITC0649, Plantain, Pome AAB) to test their response to chitosan. We have also evaluated the effect of chitosan on these cultivars by transcriptomic analysis of key genes involved in plant hormone biosynthesis, signalling and transport. We have also tested the induction of resistance in banana plants exposed to chitosan then infected with the banana wilt fungal pathogen FocTR4. This study aims to develop chitosan for preventing or managing banana root diseases sustainably.

## Results

### Chitosan depolarizes banana root cell membrane

Chitosan depolarizes plasma membrane (PM) of banana root cells, modifying its potential to half or more that measured in control cells (Fig. 1). All doses of chitosan, even the lowest one (0.1 mg·mL^-1^), cause PM depolarization achieving a less negative potential than controls. Rising chitosan concentration ten times increases PM depolarization and finally, the highest chitosan dose (2 mg·mL^-1^) causes irreversible membrane depolarization since after a recovery treatment of more than 15 minutes in the control solution, roots do not go back to non-excited membrane potential. Our findings are in the line of previous results in which authors described chitosan as a key factor to depolarize root cell plasma membrane in tomato plants^26^.

**Figure 1.**
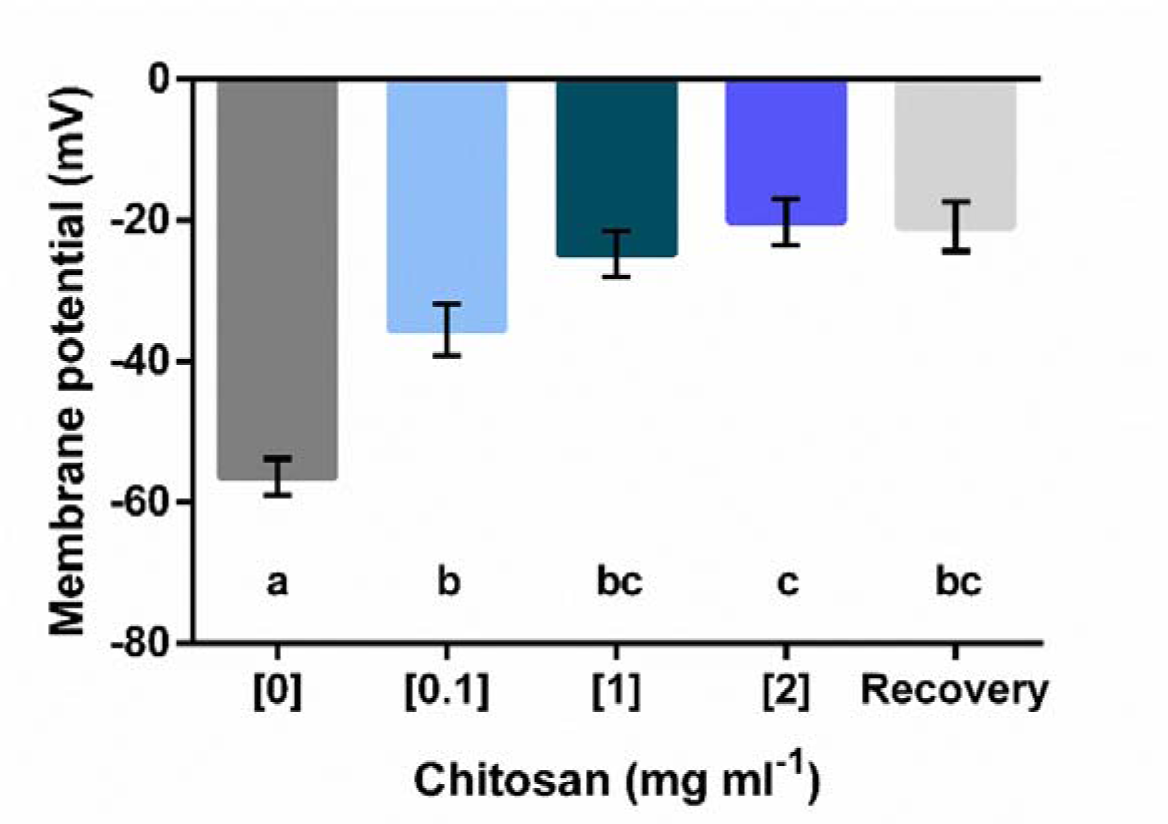
Chitosan depolarizes plasma membrane of banana root cells (cv. Petite Naine). Membrane potential variation of banana root cells treated with chitosan. Not only high (1 and 2 mg·ml ^-1^) but also low doses (0.1 mg·ml ^-1^) of chitosan decrease membrane potential of root cells when compared to the control (N=12). Membrane potential was not restored when submerging roots pre-treated with 2 mg·ml ^-1^ chitosan into the control solution for 15 minutes (Recovery; N=6). Error bars: standard error of the mean (SEM). Different letters indicate significant differences based on comparison over treatments (ANOVA and Tukey; α = 0.05).

### Chitosan reduces membrane fluidity of banana root cells

Membrane lipids of banana roots were analysed after chitosan irrigation ([0.1] and [1] mg·mL^-1^) of banana plants for 10 or 20 days (Fig. 2 and Table S1). Chitosan (1 mg·mL^-1^) induces a significant increase of myristic acid (14:0) after 10d treatment of banana roots (Fig. 2a). This trend is still present after 20 days treatment with chitosan although differences are not significant (Fig 2a). Conversely, chitosan induces a reduction in several unsaturated fatty acids (FAs) in banana roots (Fig 2b and 2c). Regarding monounsaturated FA, the percentage of eicosanoid acid (20:1n-9) decreases with high doses of chitosan (1 mg·mL^-1^) after 10- and 20-day treatments (Fig. 2b). Consequently, total monounsaturated FAs of banana membranes are reduced in chitosan-treated plants (Fig. 2b). Content of long highly polyunsaturated FA, arachidonic acid (20:4n-6), is also reduced with high doses of chitosan (Fig. 2c). This is associated with an increase of linoleic acid (18:2n-6), shorter than arachidonic acid and with less unsaturation. The observed modifications in fatty acid composition of banana roots would contribute to reduce membrane fluidity.

**Figure 2.**
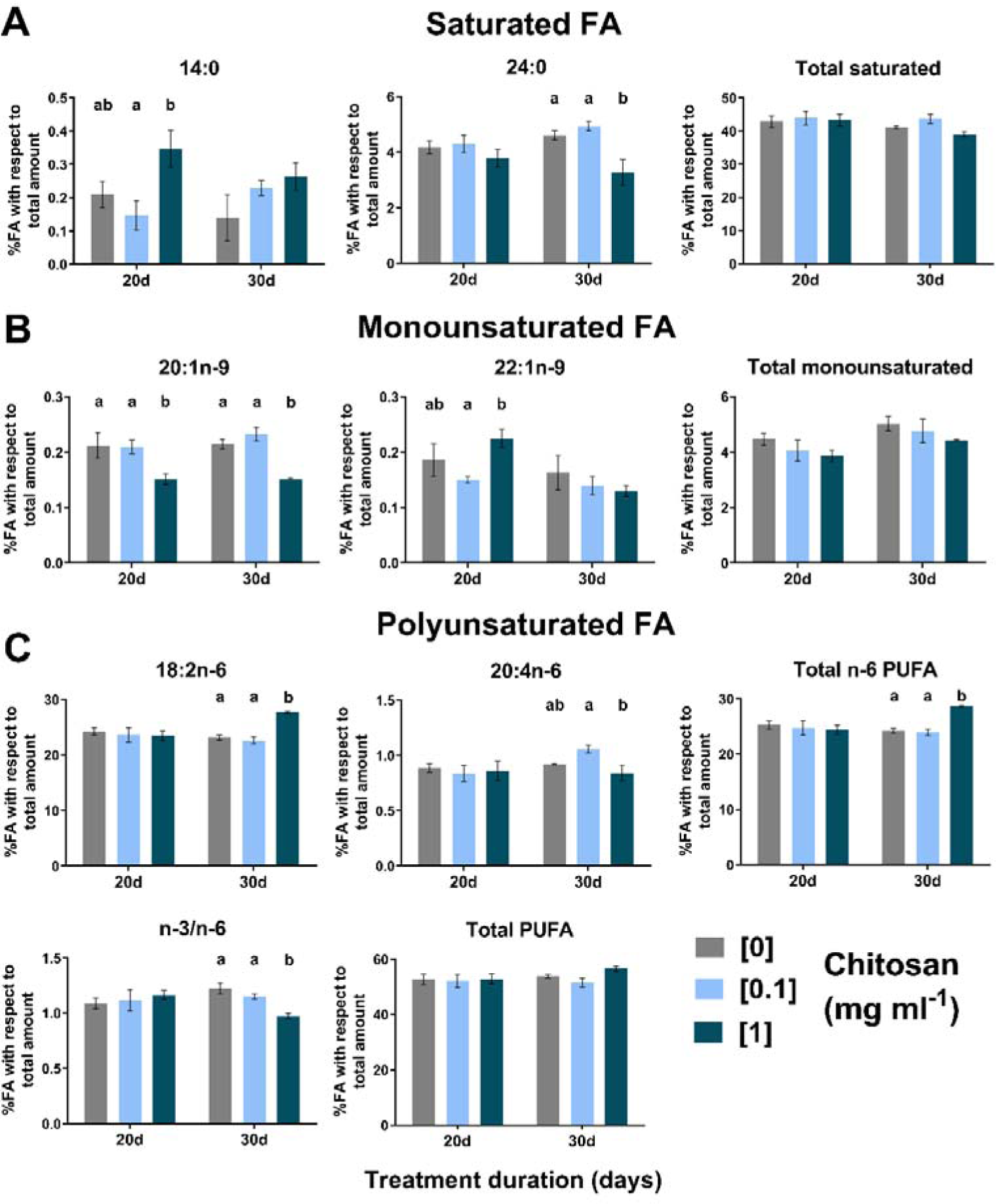
Chitosan modifies the lipid membrane composition of banana root cells (cv. Petite Naine) over time by increasing n-6 polyunsaturated fatty acids (PUFA) and decreasing saturated and monounsaturated fatty acids (FA). Membrane lipids of banana roots were analyzed in plants irrigated with different doses of chitosan for 10 and 20 days. Only FA showing significant differences among treatments are represented. **A) Saturated fatty acids (FA)**. 14:0 = Myristic acid. 24:0 = Lignoceric acid (tetracosanoic acid). **B) Monounsaturated FA**. 20:1n-9 = Eicosanoid acid. 22:1n-9 = Erucic acid. **C) Polyunsaturated FA**. 18:2n-6 = Linoleic acid. 20:4n-6 = Arachidonic acid. n-3/n-6 = ratio between n-3 and n-6 PUFA. Error bars: standard error of the mean (SEM). Different letters indicate significant differences based on comparison over treatments (ANOVA and Tukey; α = 0.05) (N treatment/control=12/12; d= days).

### Chitosan induces biosynthesis and exudation of salicylic acid and other phytohormones in banana roots

Chitosan induces stress-related phytohormones salicylic acid (SA), jasmonic acid (JA) and abscisic acid (ABA) in banana. We have detected a high amount of them in the exudates of banana roots exposed to chitosan (Fig 3a-3c). Chitosan (2 mg·mL^-1^) also induces SA and JA accumulation in banana root tissues (Fig. 3d and 3e). No differences were found in ABA in root tissues (Fig. 3f). EEM fluorescence shows dynamics of SA exudation induced by chitosan in banana roots (Fig. S1 and Table S2). Components 1 and 2 (indole-3-acetic acid, (IAA) and SA, respectively; Table S2) were the most exudated after 1, 3 and 5 days, in roots exposed to chitosan (Fig. S1a, S1b and S1c).

**Figure 3.**
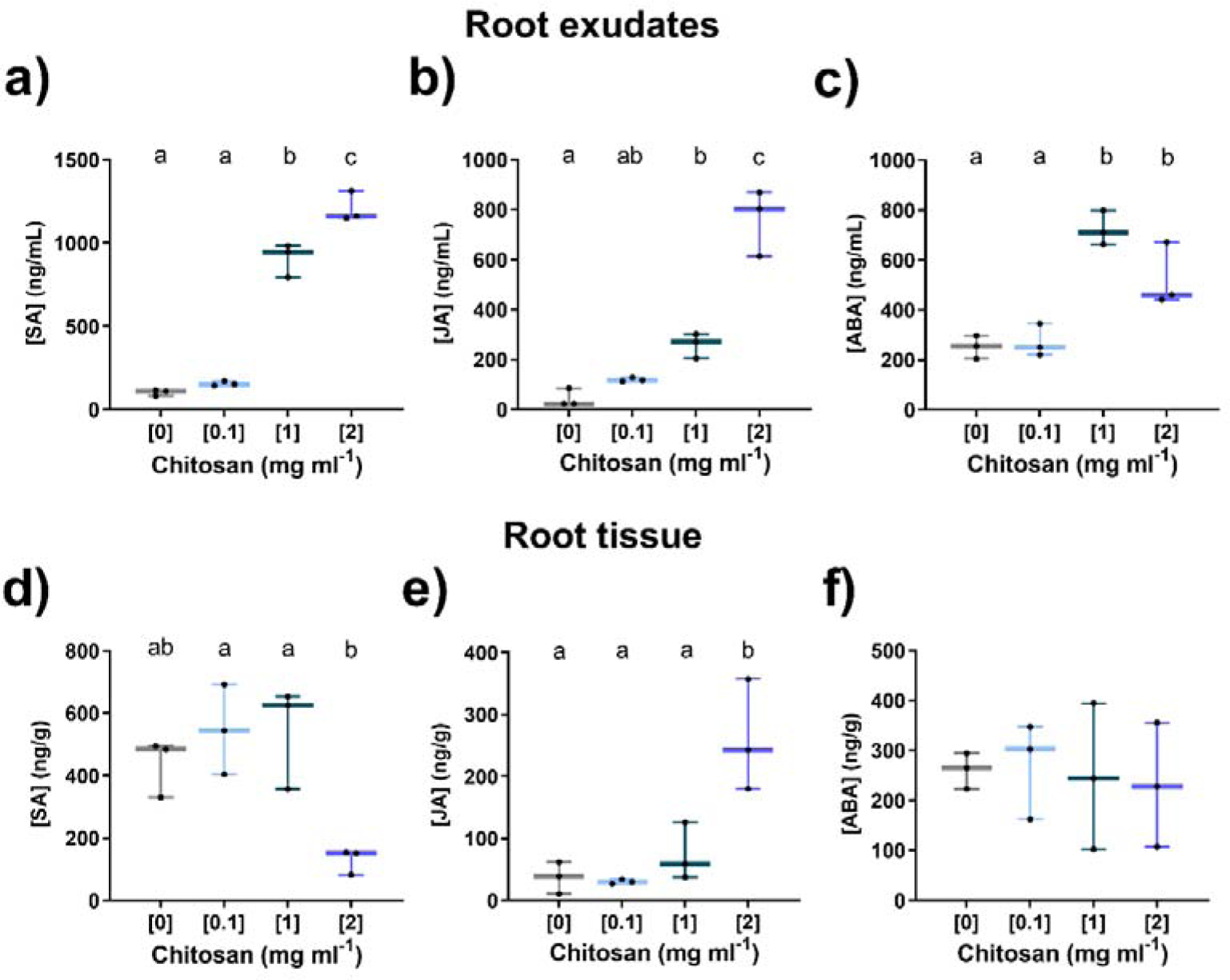
Chitosan induces exudation of salicylic acid (SA) and other phytohormones in banana roots (cv. Petite Naine) as measured by HPLC-MS. Root exudates: a) Salicylic acid (ng/g) b) Jasmonic acid (ng/g) c) Abscidic acid (ng/g). There is an increase in JA. SA and ABA levels when plants are exposed to high doses of chitosan (1 and 2 mg ml ^-1^). **Root tissue: d) Salicylic acid (ng/g) e) Jasmonic acid (ng/g) f) Abscidic acid (ng/g)**. Chitosan induces the production of JA and the accumulation of SA in banana roots (2 mg ml ^-1^). The figures correspond to the accumulation of the studied phytohormones over 3 days in three biological replicates for roots and three pools of exudates. each of three biologically independent plants. JA= jasmonic acid; SA= salicylic acid; ABA= abscisic acid. Error bars: standard error of the mean (SEM). Different letters indicate significant differences based on comparison over treatments (ANOVA and Tukey; α = 0.05).

### Chitosan increases biosynthesis of methyl salicylate (MeSA) and other plant defences volatile organic compounds (VOCs)

The hormone derivative methyl salicylate (MeSA) is induced by chitosan concentration-wise (Fig. 4a). MeSA is a volatile compound which is involved in plant physiology, and it is also described as herbivore-induced plant volatile (HIPV) known to attract the natural enemies of herbivores in agro-ecosystems^35^. Other VOCs, also involved in plant communication and defence, are induced by chitosan too. 2-Hexenal (Fig. 4b) is early induced by chitosan and perhaps originates from oxidation of membrane FA. Anisole (Fig. 4c), its derivative 3,5-dimethylanisole (Fig. 4d) and the monoterpene β-ocimene (Fig. 4e) are induced by a high (5 mg·mL^-1^) chitosan dose. 8-Methyl-heptadecane (Fig. 4f) is slightly induced by chitosan after 72h. Chitosan induce the biosynthesis of a set of VOCs which are essential in plant communication.

**Figure 4.**
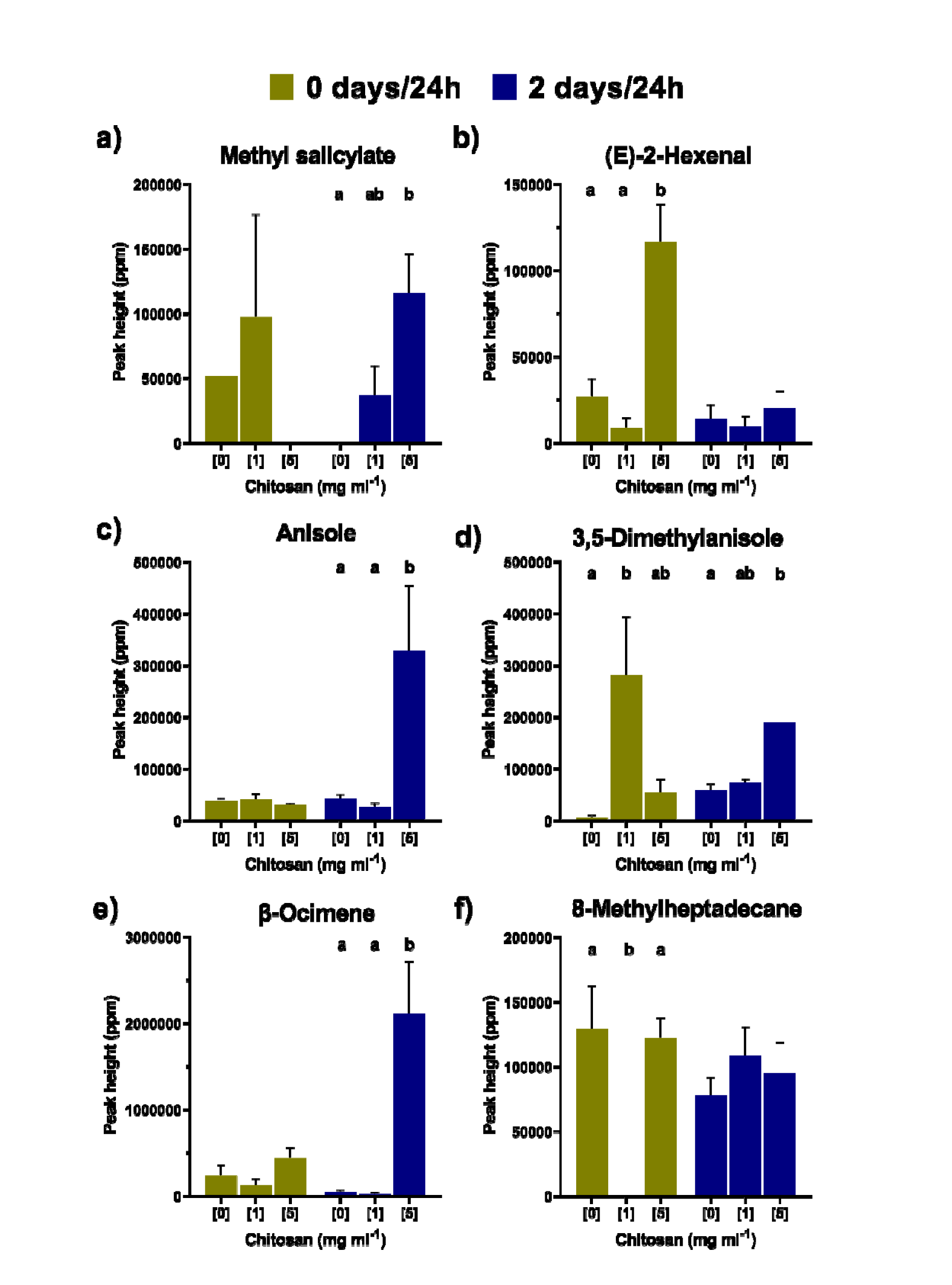
Chitosan increases production of methyl salicylate (MeSA) and other volatile organic compounds (VOCs) in banana plants (cv. Petite Naine). Chitosan increases the production of VOCs after only one day. being the differences even higher after three days. **a) Methyl salicylate b) (E)-2-Hexenal c) Anisole d) 3.5-Dimethylanisole e) β-Ocimene f) 8-Methylheptadecane**. Error bars: standard error of the mean (SEM). Different letters indicate significant differences based on comparison over treatments (ANOVA and Tukey; α = 0.05). N control/treatment = 6/3.

### Chitosan induces expression of genes involved in SA biosynthesis and systemic acquired resistance in different banana genotypes

Key banana genes related to plant defence, involved in the biosynthesis, or signalling of SA, and SA induced systemic acquired resistance (SAR), are early (1d after treatment) induced by chitosan in 60-day old plants (Figs. 5, 6, S2 and Table S3). There is a general down-regulation of gene expression 3d after treatment (Fig. 5). FMO1, involved in SA induced SAR, is the most early (1d) up-regulated gene by chitosan in all banana genotypes tested (Fig. 6). It is still up regulated (to a lesser extent) 3d after treatment in all genotypes (Fig. S2). Pathogenicity-related protein-1 (PR1) is the next most up-regulated gene by chitosan (Fig. 6). The plant immune regulator EDS1 is up regulated after 1d to a lesser extent (Figs. 6 and S2). After 3d, it is only up regulated in *Foconah* genotype (Fig. S2). Other genes such as lipase-like gene (PAD4), the specific SA receptor (NPR1), the isochorismate synthase 1 (ICS1) and phenylalanine ammonia-lyase (PAL), mainly involved in SA biosynthesis, are also up regulated by chitosan 1d after treatment (Fig. 6). This situation is reversed 3d after treatment (Fig. S2). However, the polyphenol oxidase (PPO) is highly repressed at both times.

**Figure 5.**
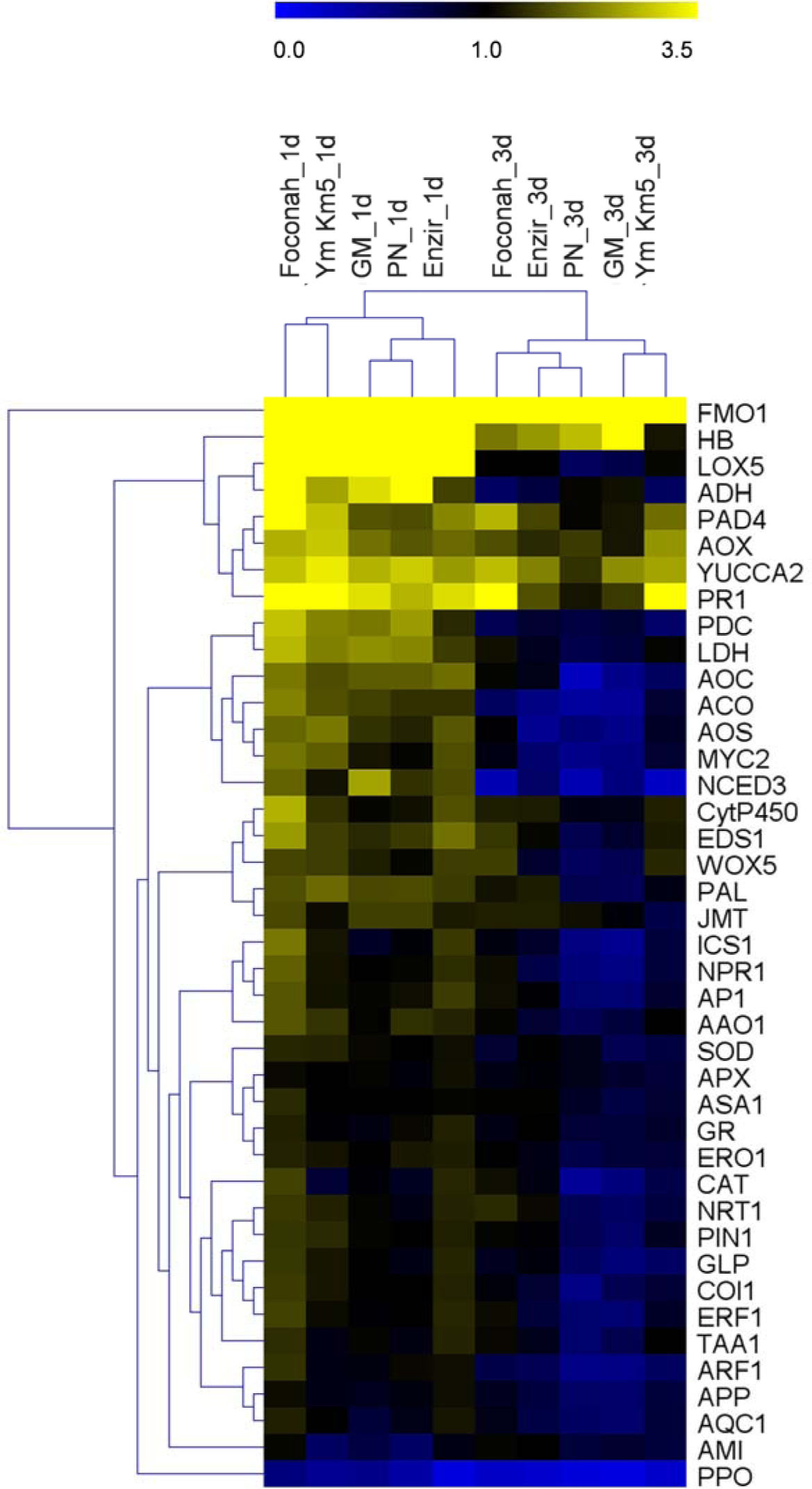
Effect of chitosan on differential expression of defense growth and hypoxia genes in five banana cultivars. Two-way hierarchical clustering of the differential expression for 42 genes quantified in roots of five banana genotypes treated with chitosan in hydroponic tray system. Manhattan distance and average linkage clustering were used to group the analyzed genes and genotypes. Each cell represents the average fold change (FC) of 6 independent biological replicates in each genotype and time point and is relative to the mean control. *Musa genes* EF-1 and *L2* were used as internal controls to normalize the expression data. Black: no differential expression (FC=1). yellow: up-regulation (FC>1). blue: down-regulation (FC<1). Enzir: Enzirabahima; GM: Gros Michel; PN: Petite Naine; Ym Km5: Yangambi Km5. d: day(s) of chitosan treatment (1 mg ml^-1^*). N control/treatment=6/6*.

**Figure 6.**
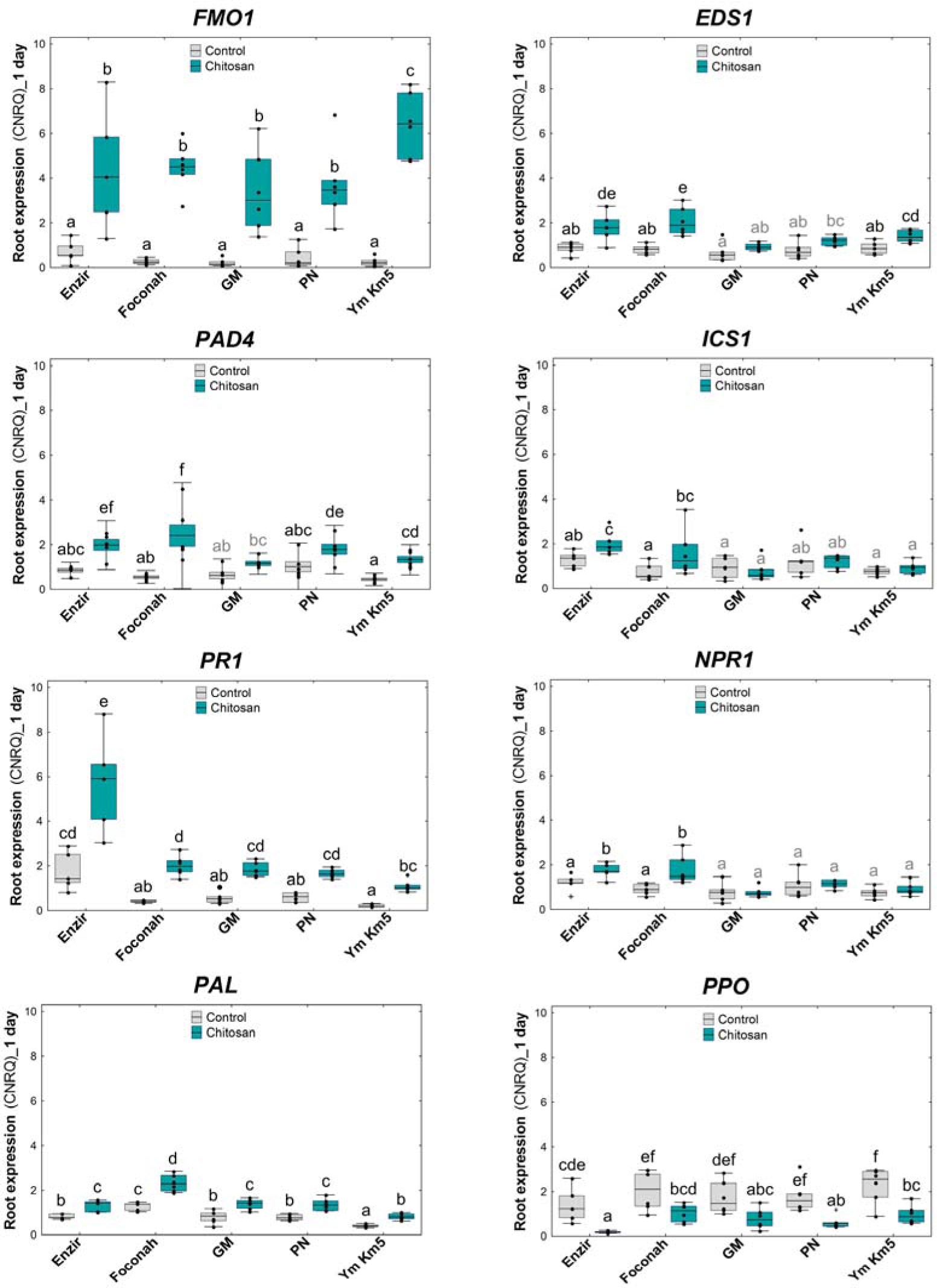
Chitosan induces expression of genes involved in salicylic acid (SA) metabolism and SAR in five banana genotypes exposed 1 day to chitosan. Relative gene expression levels quantified in banana roots of diverse genotypes subjected to chitosan treatment in hydroponic tray system. CNRQ: Calibrated Normalized Relative Quantity. *Musa genes EF-1* and *ACT-1* were used as internal controls to normalize the expression data. Error bars = standard error of the mean (SEM). Different letters indicate significant differences based on comparison over treatments. genotypes and genotype x treatment (Two-way ANOVA and Tukey; α = 0.05). N control/treatment= 6/6. Gene abbreviations according to Table S3.

### Chitosan induces genes involved in JA metabolism and oxidative status of the cell

Allene oxide cyclase (AOC), allene oxide synthase (AOS) and the S-adenosyl-L-methionine:jasmonic acid carboxyl methyltransferase (JMT) genes, involved in JA biosynthesis and signalling pathways, are induced after 1d chitosan treatment (Figs. 5 and S3). JMT is still up regulated after 3d (Fig. S3). Super oxide dismutase (SOD) and endoplasmic reticulum oxidoreductase 1 (ERO1) genes related to ROS are induced after 1d chitosan treatment (Fig. S3). After 3d, all the chitosan treatments show ROS genes down regulation in all banana genotypes. However, all hypoxia-related genes tested (HB, ADH, AOX, PDC and LDH) are early induced by chitosan (Fig. S3). Genes related to parasitism and endophytism (mainly CytP450 and to a lesser extent AP1 and NRT1) are induced by chitosan after 1d treatment (Fig. S3).

### Genes related to banana plant growth are induced by chitosan

Expression of indole-3-acetaldehyde oxidase (AAO1) and the auxin efflux carrier PIN1 genes, but especially YUCCA2, a flavin-containing monooxygenases which catalyse the conversion of indole-3-pyruvic acid (IPA) to IAA, related to auxin biosynthesis and mobilisation are up-regulated after 1d chitosan treatments in banana roots (Fig. 5 and Fig. S3). Expression of banana genes related to ethylene and ABA biosynthesis are up regulated after 1d of exposure to chitosan, however, these genes are repressed in banana plants exposed 3d to chitosan (Fig. 5 and Fig. S3). Root growth and development related genes, lipoxygenase-5 (LOX5) and WOX5, a member of the Wuschel (WUS) family of homeodomain transcription factor, both are induced with 1d-chitosan treatment in banana roots (Figs. 5 and S3).

### Banana cultivars differ in their transcriptomic response to chitosan

We tested the effect of banana genotype on gene expression modifications by chitosan, using a subset of five banana cultivars namely: *Petit Naine*, *Gros Michel*, *Enzirabahima*, *Yangambi Km5* and *Foconah*. They cover most of the genetic diversity of the banana crop available (Fig. S4). Banana cultivars differ in their transcriptomic response to chitosan. Banana cultivars with resistance to both, FocR1 and TR4 (or STR4), *Enzirabahima*, *Yangambi Km5* and *Foconah* have the greatest number of genes up-regulated after 1 and even 3-day chitosan treatments (Fig. 5). Chitosan causes one day after treatment in *Yangambi Km5* for FMO1 gene, the largest induction of all genotypes tested (ca. 7-fold, Fig6). For PR1 gene, chitosan causes a ca. 6-fold induction in *Enzirabahima* genotype, significantly larger than the rest. Both genes are involved in the establishment of SAR and related to the presence of SA.

### Chitosan induces resistance to the wilt pathogen FocTR4 in banana

We have tested the effect of chitosan, the endophytic and chitosan-resistant biocontrol agent *P. chlamydosporia* strain 123 (Pc123)^14^ and both together on the colonization of roots from healthy banana plantlet roots by *F. oxysporum* f sp. *cubense* Tropical Race 4 (FocTR4). This wilt fungus is the causing agent of the serious new outbreak of banana *Panama Disease* spreading worldwide. Treatments were deployed prior to inoculation (pretreatment) with the wilt pathogen. All treatments significantly reduce FocTR4 colonization of banana roots respect to untreated controls (Fig. 7a). Plantlets treated with either Pc123 or “Pc123 + chitosan” show the least (18%) total root colonization by the pathogen. Chitosan only reduces to 40% FocTR4 colonization of roots. Finally untreated controls show 65% root fragments colonized by FocTR4. Treatments have a similar effect reducing FocTR4 endophytic colonization of banana roots (Fig. S5). Real-time PCR confirms the results of culturing (Figs. 7b and S6). However, the molecular method is more sensitive than culturing and shows that treatments with chitosan (alone or with Pc123) are more effective reducing FocTR4 banana roots colonization that Pc123 on its own.

**Figure 7.**
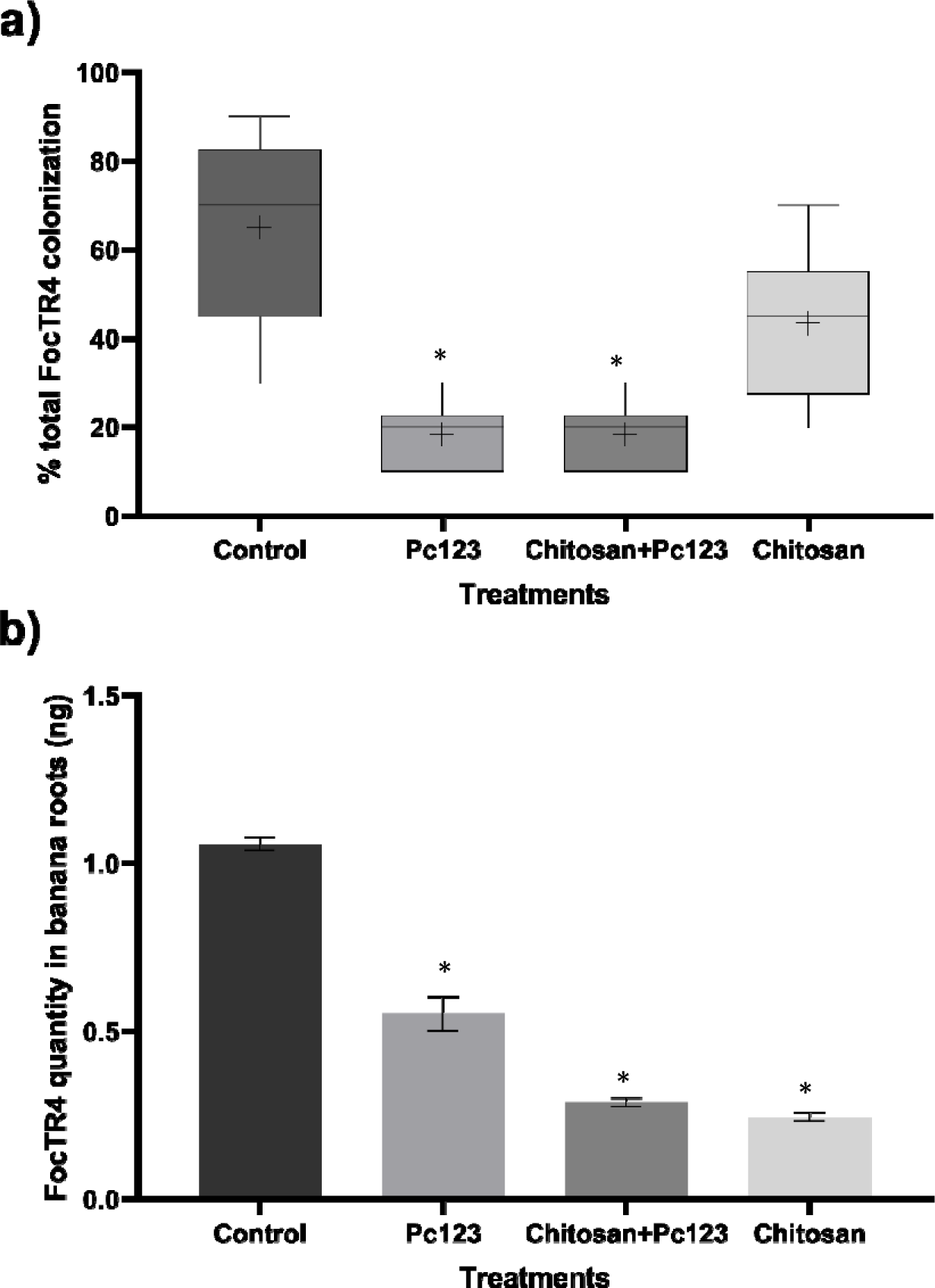
Chitosan and the biocontrol fungus *P. chlamydosporia* induces banana resistance to Fusarium wilt. **a)** Quantification of *P. chlamydosporia* strain 123 and *Fusarium oxysporum* f. sp. *cubense* TR4 total root colonizationby culturing techniques. **b)** RT-PCR quantification of FocTR4 root colonization. * Indicates significant differences (P-values <0.05)

### Chitosan promotes growth of *Musa* spp. plants under greenhouse conditions

Chitosan irrigation (0.1 and 1 mg·mL^-1^) for 30 days does not affect banana plantlet growth (Fig. 8a). A larger dose (2 mg·mL^-1^) of chitosan significantly reduces root and shoot growth as described for Arabidopsis ^12^. Long term irrigation (70 days) in the greenhouse with 1 mg·mL^-1^ of chitosan significantly promotes pseudo stem and corm growth (length) (Fig. 8b). There is also a slight promotion of the root weight.

**Figure 8.**
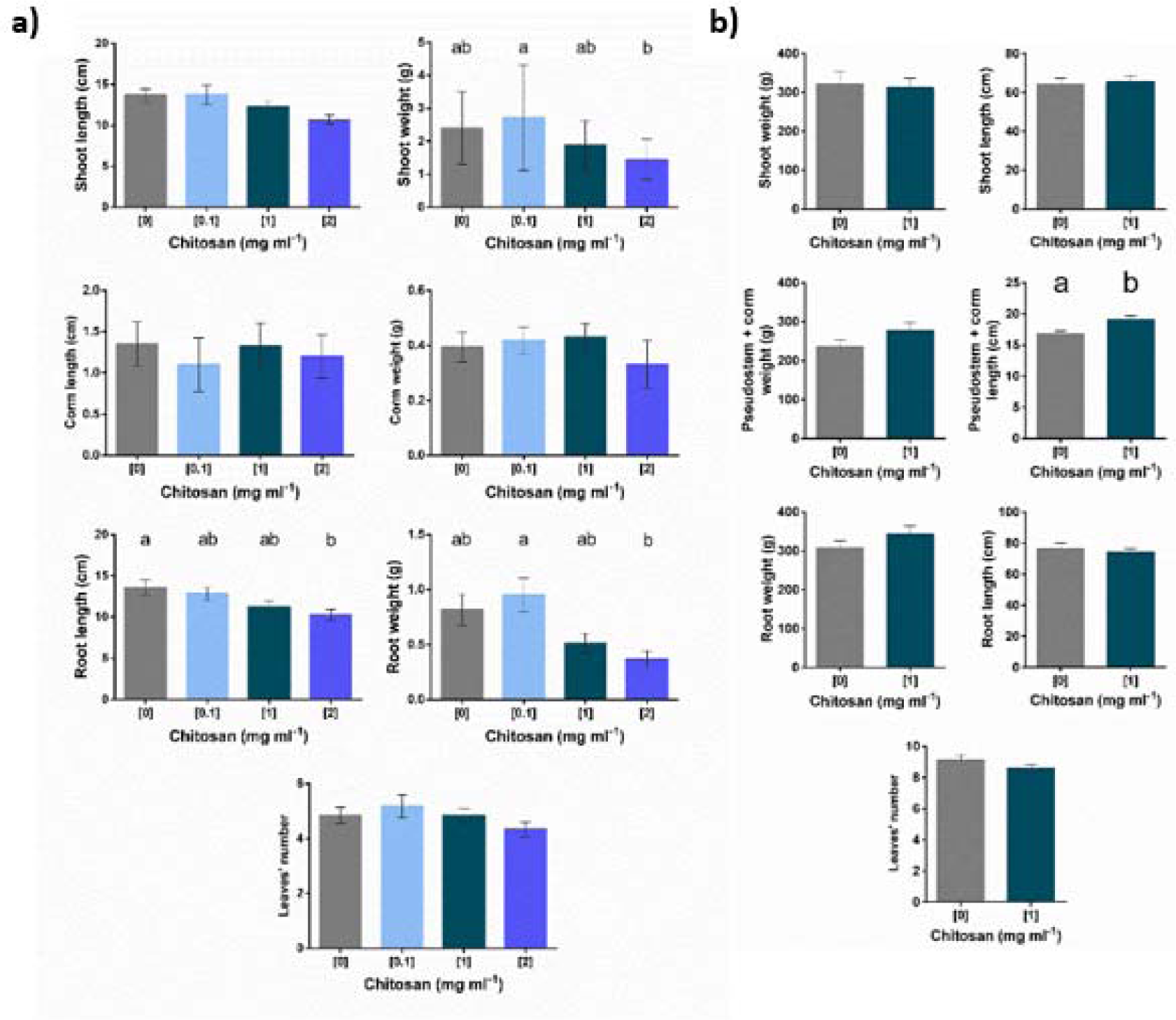
Chitosan irrigation alters the growth of banana plants (cv. Petite Naine). **a) 30-days irrigated pot plants.** Irrigation at the highest dose (2 mg·mL ^-1^) inhibits banana root and shoot development. N= 15. Error bars = standard error of the mean (SEM). Different letters indicate significant differences based on comparison over treatments (ANOVA and Tukey; α = 0.05). **b) Greenhouse plants**. There is a growth promotion effect in the plants irrigated with 1 mg/mL dose of chitosan for 10 weeks. Error bars = standard error of the mean (SEM). N= 15. Different letters indicate significant differences based on comparison over treatments (ANOVA and Tukey; α = 0.05).

## Discussion

We show that chitosan induces defences in banana plants (*Musa acuminata*) which reduce root colonization by FocTR4, a serious banana wilt pathogen ^47^. Chitosan is a plant defence elicitor^25^ with antimicrobial activity against plant pathogenic bacteria, fungi, and nematodes^13,14,25,48^.

We give evidence that chitosan applied in the rhizosphere induces genes involved in banana SA metabolism. Chitosan modulates immune responses^49,26^ and enhances activities of beneficial microorganisms naturally present or added ^21,50,51,52^.

In this work, we show that chitosan depolarizes banana root cell plasma membrane to a higher extent than for tomato plants^26^. Plasma membrane is a well-established chitosan target for fungi and plants^53,54,55^. Chitosan has also been proposed as a potential transdermal penetration enhancer in mice, decreasing cell membrane potential (HaCaT cell line) and enhancing cell membrane fluidity^56^.

When chitosan depolarizes/permeabilizes the cell membrane triggers an increase in reactive oxygen species (ROS)^55^ and secondary metabolites^57^ in fungi. A ROS burst is also evident in *Arabidopsis* plant roots exposed to chitosan^12^. In our study, chitosan induces ROS-related genes (SOD and ERO1) in banana. Banana cultivar *Petite Naine* adapts to chitosan stress by reducing membrane fluidity increasing myristic acid (14:0) and decreasing eicosanoid (20:1n-9) and arachidonic (20:4-6) acids amounts. A fatty acid desaturase mutant (Δods) of the model fungus *Neurospora crassa* with reduced plasma membrane fluidity showed increased resistance to chitosan in comparison with the wild-type^53^.

We have found a significantly increase in the content of JA, a lipid derived phytohormone, in banana root tissue and exudates of chitosan treated plants. Accordingly, JA biosynthesis genes AOC, AOS and JMT^58,59^ are early overexpressed. The oxylipin oxophytodienoic acid (OPDA)^60^ is a precursor of JA. Oxylipins are produced from long chain fatty acids^61^, so they may be used for JA production or released to the rhizosphere. To this respect, we have detected 2-(E)-hexenal in the volatile profile of chitosan primed banana plants. This aldehyde occurs in stressed plants, and it could be derived from oxidation of membrane FAs, such as linoleic acid^62,63^.

Auxin (AAO1, PIN1 and YUCCA2) and transcription factors related to root cell differentiation in the plant quiescent centre (LOX5 and WOX5) genes are induced in chitosan-treated banana roots. Therefore, chitosan irrigation (up to 1 mg mL^-1^) promotes growth of 70d-old banana plants. However, a larger chitosan dose (2 mg mL^-1^) reduces growth as found for Arabidopsis^12^.

Chitosan induces expression of SA biosynthesis and signalling (PR1, EDS1, PAD4, NPR1, ICS1 and PAL) and SA induced systemic acquired resistance (FMO1) genes in banana plants. In immunity related processes, EDS1 and PAD4 genes promote SA accumulation^64^. As a result, chitosan induces root exudation of SA and systemic production of MeSA and other defence volatiles in banana plants. This could be due to the role of chitosan as mild pathogen/microbe-associated molecular pattern (PAMP/MAMP)^65^. Hormone changes in the roots and exudates of banana plants would be related to the systemic effect of chitosan in the plant. It seems that SA its rapidly synthesized and exudated to the rhizosphere since it is not accumulated in banana root tissue. Alternatively, SA accumulation could be produced sooner than 3 days which was the time we established to collect the samples. JA is only accumulated in roots tissue and secreted (exudates) with highest chitosan dose (2 mg·mL^-1^). Chitosan also causes JA and SA accumulation in Arabidopsis root tissues^12^. Foliar application of chitosan also induces JA and SA plant hormones stress responses ^59, 66^. Moreover, SA induces synthesis and accumulation of phenols also reflected in our EEM fluorescence analysis ^67^.

*Petit Naine*, *Gros Michel*, *Enzirabahima*, *Yangambi Km5* and *Foconah* banana cultivars differ in their responses to chitosan. Banana genotypes display diverse transcriptomic profiles in reply to a mild osmotic stress^68^. Chitosan causes in Foc Race 1-resistant, *Yangambi Km5* ^69^ the largest induction (ca. 7-fold) of FMO1 gene of all genotypes tested. FMO1 is an essential component of SAR in Arabidopsis^70^. Chitosan induction of SA (local and systemically) is reflected in all steps of SA biology. Both SA, isochorismate (IC) and phenylalanine ammonia-lyase (PAL), biosynthetic pathways^71^ encoding genes (ICS1 and PAL) are induced by chitosan in banana plants as described also in Arabidopsis^12^. Then, SA receptor encoding gene NPR1^72^ is significantly induced by chitosan in *Enzirabahima* and *Foconah* banana genotypes. Chitosan causes a ca. 6-fold induction of the SA induced response gene PR1^27^ in *Enzirabahima* genotype, significantly larger than the rest. EDS1 and PAD4 genes involved in effector triggered immunity (ETI) and basal immunity^73^ are induced significantly by chitosan in most banana genotypes tested. These priming results are very relevant since banana genotypes show limited resistance to FocTR4^74^. Future studies should test chitosan defence induction in banana hybrids resistant to FocTR4^75^.

Our transcriptomic responses of banana to chitosan are reflected in chitosan primed banana response to pathogens. Preventive treatments of chitosan with the endophytic biocontrol agent Pc123 reduce (ca. 4-fold) colonization of banana roots by the wilt pathogen FocTR4. FocTR4 is causing a serious new outbreak of banana wilt which is quickly spreading^76^. To this respect FocTR4 biocontrol approaches have been carried out with non-pathogenic *Fusarium oxysporum, Trichoderma* spp, *Serratia* spp. and *Burkholderia* spp and *Streptomyces* sp.^77,78,79^. Therefore, chitosan alone or in combination with compatible beneficial endophytes can help manage sustainably FocTR4, and perhaps other biotic stresses, in banana. Previous studies described the link between biocontrol agents (BCA, fungi and bacteria) with a plant induction of SA and SAR. For instance, the BCA *Pseudomonas simiae* PICF7 in banana plants modifies the expression of defense-related genes in banana roots. *P. simiae* initially suppresses SAR and induced systemic resistance (ISR) related genes trying to avoid the banana immune response^80^. This bacterium was also evaluated in combination with Foc Sub Tropical Race 4 and decreased the severity of Fusarium wilt^81^, may be by inducing these defence-related genes. Its combination with chitosan could improve their efficacy under field conditions and, the reduction of the fungus in the plant, as it is known that chitosan affects Foc sporulation, mycelial growth, and germination^82^. *Trichoderma harzianum* T34 also induces an early repression of JA and SA-defence related genes in *A. thaliana* after inoculation with the fungus^83^. Previous work from our laboratory also described the biocontrol agent *P. chlamydosporia* and chitosan as inducer of plant defence hormones pathways^12,84^. However, this work is the first report in which it is described the effectiveness of the combined use of the biocontrol fungi *P. chlamydosporia* and chitosan applied together. We demonstrate that both, *P. chlamydosporia* and chitosan, are complementary to prime banana plants and protect them against banana wilt FocTR4. In this line, recent studies have described that biocontrol fungi are able to induce secondary metabolisms and overproduce VOCs under their exposure to chitosan. One of the most emitted VOC by *P. chlamydosporia* when it grew with chitosan is oct-1-en-3-ol^85^. This volatile is described as inhibitory compound of plant pathogens such as *Penicillium paeum* or *Aspergillus flavus* ^86,87^. In addition, the increases of the emission of MeSA by banana plants treated with chitosan can also act as natural repellent for *Cosmopolites sordidus,* the main pest threatening banana plantations. We, therefore, have demonstrated that chitosan can be used to prime banana defences local and systemically, and as it is able to induce banana growth regulators, it has a fertilizer role too, as proved in other studies^88,89^. Therefore, this polymer could be used against FocTR4. Chitosan irrigation could also be a tool for managing above ground banana pests and diseases since it induces SAR compounds such as MeSA.

## Material and methods

### Plant material, chitosan preparation and experimental system

Five triploid genotypes (AAA) representing important cultivated subgroups of bananas were used in this study: *Petite Naine* or Dwarf Cavendish (subgroup Cavendish), *Gros Michel* (subgroup Gros Michel, ITC1122), *Enzirabahima* (subgroup Mutika/Lujugira, ITC1354), *Yangambi Km5* (subgroup Ibota Bota, ITC1123), and *Foconah* (subgroup Pome/Prata, ITC0649) (Table 1). *I n v i t r*p*o*ropagated plants of the genotype *Petite Naine* were kindly provided by CULTESA (Cultivos y Tecnología Agraria de Tenerife S.A, Tacoronte, Sta. Cruz de Tenerife, Spain), while plants of the genotypes *Gros Michel, Enzirabahima, Yangambi Km5* and *Foconah* were obtained through the International Musa Transit Center (ITC, Bioversity International) hosted at KU Leuven, Belgium.

**Table 1.** Description and characteristics of the five Musa genotypes evaluated in this study.

| Name | <sup>a</sup> ITC Code | Ploidy Level | Genomic constitution | Type | Subgroup | Source/origin | Distribution | *Foc* response |
| --- | --- | --- | --- | --- | --- | --- | --- | --- |
| “Petite Naine”<br>(Dwarf Cavendish) | NA | 3x | AAA | Dessert | Cavendish | <sup>b</sup> CULTESA/Canary Islands (Spain) | All continents: (sub)tropical areas | *Foc* R1 resistant;<br>*Foc* (S)TR4 susceptible <sup>1</sup> |
| “Gros Michel” | ITC1122 | 3x | AAA | Dessert | Gros Michel | <sup>c</sup> CIRAD collection in Guadeloupe/unknown | All continents: (sub)tropical areas | *Foc* R1 and TR4 susceptible <sup>2</sup> |
| “Enzirabahima” | ITC1354 | 3x | AAA | Beer and cooking | Mutika/Lujugira | <sup>d</sup> KARI/Uganda | East Africa. Colombia | *Foc* R1 and TR4 resistant <sup>3</sup> |
| “Yangambi Km5” | ITC1123 | 3x | AAA | Dessert | Ibota Bota | <sup>c</sup> CIRAD collection in Guadeloupe/Democratic Republic of Congo | Africa. Indonesia. Thailand | *Foc* R1 and STR4 resistant <sup>4,5</sup> |
| “Foconah” | ITC0649 | 3x | AAB | Dessert. sweet | Pome/Prata | <sup>c</sup> CIRAD collection in Guadeloupe/Cameroon | West Africa. India. Malaysia. | *Foc* R1 and TR4 resistant <sup>6</sup> |

Agricultural Research Institute. NA: not available. *Foc*: *Fusarium oxysporum* f.sp.*cubense*; R1: Race 1; TR4: Tropical Race 4. STR4: Subtropical Race 4.

Medium-size molecular weight chitosan (70 kDa) with 90.1% deacetylation degree (T8s) (Marine BioProducts GmbH, Bremerhaven, Germany) was prepared^14^ and used in all experiments as liquid solution. To study the effect of chitosan in banana roots at the molecular or physiological level, a tray system experimental set-up was used. Six L-polypropylene containers were used, and chitosan was applied to the hydroponic medium^90^, with the exception that 112 μL L^-1^ of Fe-EDDHSA was added instead of sequestrene. Each tray was filled with 3.5L of medium and connected to an air pump to keep oxygen levels under normoxia ([O_2_] = 7-8 mg L^-1^). Trays were covered with lids with holes and five banana plants (one of each genotype) were randomly distributed in each tray with their roots submerged into the hydroponic medium (Supplementary Fig. S6). Plants (12 replicates per genotype) were grown for 30 days under controlled conditions: 12 h/12 h (light/dark cycle), 25°C, 85% RH and 65 µmol m^-2^s^-1^ light intensity at leaf surface (on average). During this period, the medium was renewed weekly and dissolved oxygen was measured twice per week with an optical oxygen sensor probe (Firesting O_2_, Pyroscience). Subsequently, the assay was started by adding fresh medium with chitosan at 1 mg·mL^-1^ to half of the trays (treated plants) while the other half received fresh medium without chitosan (control plants).

### Root electrophysiology experiments

Excised banana roots were placed individually in a holder chamber filled with 1/10 Gamborg’s nutrient solution B5 1:10 (Basal Medium Minimal Organic, Sigma^91^) (control) and the experiment was carried in growth chambers^26^. Glass microcapillaries filled with 0.5M KCl (tip diameter < 1 µm) were inserted into root cortical cells until a stable potential was obtained ^92^. This initial membrane potential was recorded and afterwards roots were exposed to increasing concentrations of chitosan (0.1, 1, and 2 mg mL^-1^) in Gamborg’s 1/10 solution and for each treatment membrane potentials were recorded once a stable reading was obtained. The experiment was incremental and between each treatment with chitosan root medium as replaced with control solution and kept for 1 minute to allow the root to recover. After the highest dose of chitosan, the bathing solution was changed again with fresh control Gamborg’s 1:10 solution, and the roots were allowed to recover for a longer period (15 min) and the final membrane potential was recorded.

Twelve biological replicates per treatment were used for the experiments and the statistical study of the data obtained was performed using JMP Pro 13.2.1. The normality of the samples was checked using the Shapiro-Wilk Normality test and Levene’s test to check the homogeneity of variances. ANOVA was performed using TukeyHD test, (α = 0.05)^93^.

### Membrane lipid composition of fatty acids

Banana plants were placed individually in pots with sterile peat and amended for 10 days with Gamborg’s B5 1:10 solution, after that, plants were irrigated with different solutions of chitosan (0.1 and 1 mg·mL^-1^) for 10 and 20 days (20 and 30 days of total growth). Plant development parameters were measured, and the roots were frozen in liquid nitrogen. Three replicates of each freeze-dried banana roots plantlets were analysed for fatty acid analysis. Fatty acids were extracted from 0.3 to 1.0 grams of tissue samples by homogenization in 20 ml of chloroform/methanol (2:1 v/v) in an Ultra Turrax tissue disrupter (IKA ULTRA-TURRAX T 25 digital, IKA-WERKE). The total lipids were prepared^94^ and non-lipid impurities were removed by washing with 0.88% (w/v) KCl. The weight of lipids was determined gravimetrically after evaporation of the solvent and overnight desiccation in vacuum. Fatty acid methyl esters (FAME) were prepared by acid-catalysed transesterification of total lipids^95^, and the total lipid samples were transmethylated overnight in 2 ml of 1% sulphury acid in methanol (plus 1 ml of toluene to dissolve neutral lipids) at 50°C. The methyl esters were extracted twice in 5 ml hexane-diethyl ether (1:1, v/v) after neutralization with 2 ml of 2% KHCO_3_, dried under nitrogen and redissolved in 0.1 ml of iso-hexane. Methyl esters were purified by TLC (thin layer chromatography) using iso-hexane:diethyl-ether:acetic acid (90:10:1 v/v/v) and stored at −80°C till analysis. FAME were separated and quantified by gas-liquid chromatography using an SP™ 2560 flexible fused silica capillary column (100 m long, internal diameter of 0.25 mm and film thickness of 0.20 mm, SUPELCO) in a Hewlett-Packard 5890 gas chromatograph. The oven temperature of the gas chromatograph was programmed for 5 min at an initial temperature of 140°C, and increased at a rate of 3°C/min to 230°C, further increased at a rate of 2°C·min^-1^ to 240°C and then held at that temperature for 12 min. The injector and flame ionization detector were set at 260°C. Helium was used as carrier gas at a pressure of 300 KPa, and peaks were identified by comparing their retention times with appropriate FAME standards purchased from the Sigma Chemical Company (St Louis, MO, USA). Individual fatty acid concentrations were expressed as percentages of the total content. The results are expressed as mean ± standard error. Individual fatty acids data were statistically analysed by two-way analysis of variance (ANOVA) to determine differences between distance treatments. Statistical analyses were conducted using SPSS Statistical Software System version 15.0 (SPSS Inc., Chicago, IL). As fatty acid data were percentages, they were transformed with arcosen (x + 1), and all statistical tests were performed with a significance level of α = 0.05.

### Analysis of volatile organic compounds from banana plantlets

Magenta Box (MB, Sigma) halves joined one on top of the other with a gasket were used as experimental system. Five *Musa acuminata* (cv *Petit Naine*) *in vitro* culture plants were placed in each MB. Plants with similar size and number of roots were selected for experiments. The treatments for these experiments were control solution of Gamborg’s B5 1:10 medium and 1 and 5 mg·mL^-1^ chitosan diluted in the same Gamborg’s solution. We used 45 mL of the hydroponic solutions per MB. Three replicates were prepared, for the two times studied. The treatments were evaluated in two times 0 days/24h, for early volatiles, where the fibers were capturing volatiles for the first 24 hours; and 2 days/24h or late volatiles, in this case, the plants were kept in the Magenta Box with the treatments for 2 days and after that, the fibers were introduced to capture volatiles for 24 hours. Plants were incubated in a growth chamber (SANYO Plant Growth Chambers Sanyo MLR-350H), at 24°C, 60% relative humidity and with a photoperiod of 16:18 hours of light and darkness, during each experiment. We made the same measurements without banana plants, to evaluate VOCs from the hydroponic solutions alone and to assess environmental contaminants. These VOCs were later removed from the analysis. We then consider plant VOCs only.

The VOCs profile of the plants was determined by Gas Chromatography-Mass Spectrometry (GC/MS) using head space-solid phase microextraction (HS-SPME). The analysis was performed by a low-resolution mass spectrometer (Agilent model 5977B) with quad-coupled analyser coupled to a gas chromatograph (Agilent model 7890B), equipped with a capillary column with split/split less injector. Ionization of the samples is performed in electronic impact. SPME manual holders (SUPELCO) and fibers (SPME Fiber Assembly 50/30um DVB/CAR/PDMS, SUPELCO) were used for microextraction. The holder is a syringe-like mechanism with a needle passing through the mechanism and containing a delicate retractable SPME fiber that can be replaced^96^. The holder was placed over a hole previously made to the MB, so that the fiber remained inside the MB. The fiber was kept inside the container in contact with the plant environment for 24h. Then it was collected for desorption in the injector for 4 minutes. Chromatographic separation was carried out for 41 minutes. The injector was at 250°C, reaching a maximum of 260°C. The MS was run in scan mode. Finally, compound identification was performed by comparing the experimental mass spectra with those of the system library (Wiley275).

Once VOCs profiles of samples were obtained, data treatment was performed prior to the analysis of the results. The data relating to the replicates of each treatment were analysed jointly, comparing the mass spectra to verify that the same compound appeared in all the replicates. The experimental spectra were compared with those appearing in different databases, such as NIST: National Institute of Standards and Technology (https://webbook.nist.gov/chemistry/name-ser/) and PubChem Compound (https://www.ncbi.nlm.nih.gov/pccompound), assigning to each spectrum the compound with which it showed the greatest similarity. Of the compounds identified, those that only appeared in one replicate were discarded if the peak height was less than 50,000 ppm. From the VOCs identified in the plant samples, those that appeared in the hydroponic solutions were eliminated. Finally, only compounds generated by banana plants in the presence or absence of chitosan were selected. The compounds selected were classified by molecule type, using PubChem Compound (https://www.ncbi.nlm.nih.gov/pccompound) and ChemSpider (http://www.chemspider.com/) databases.

An unbalanced one-factor ANOVA was performed to detect statistical differences in the abundance, i.e., in the height of peaks (ppm) of the compounds appearing in common between samples with and without chitosan, the same procedure was performed for the two times, determining whether the differences in abundance appeared in an early phase, a later phase, or both. Subsequently, an a posteriori test, TukeyHD test, was performed with two-by-two comparisons between treatments to observe between which groups the differences were, if any. Previously, normality was checked with the Kolmogorov-Smirnov test and homoscedasticity, using Bartlett’s test, fulfilling all the requirements. When values did not fit a normal distribution, the Kruskal-Wallis non-parametric test was performed, and if differences existed, the Dunn’s test was used. The graphs and analysis of this work were performed using GraphPad Prism 8.0.1 software.

### UHPLC hormone analysis and Emission Excitation Matrix (EEM) fluorescence analysis

Banana plantlets were placed as in the experiment explained above. Each Box was filled with 50 mL of sterilized Gamborg’s B5 basal mixture 1:10, amended with chitosan at different concentrations (0, 0.1, 1 and 2 mg·mL^-1^). Plants were then placed in a growth chamber at 24°C, 60% RH with a 16:8 hours light: dark photoperiod and incubated for 3 days. This experiment was performed with ten biological replicates per treatment. Root exudates collected in this experiment are the results of 3 days accumulation. Plantlets were removed from the MB, keeping the roots for further analysis, the exudates were collected and filtered as explained before. The exudates of three biological replicates (2 mL) were mixed to obtain 3 pools per treatment of 6 mL each. Roots and pools of roots exudates, were frozen in liquid N2, lyophilized, and stored at −20°C until use. The samples were analysed for plant hormone content, specifically: salicylic acid (SA), jasmonic acid (JA) and abscisic acid (ABA) by ultra-high performance liquid chromatography-mass spectrometry (UHPLC-MS, Q-Exactive, ThermoFisher Scientific)^26^.

Banana plantlets were placed individually in sterile 575 mL Magenta Boxes (GA-7, Sigma). Each box was filled with different treatments consisting of 50 mL of sterilized Gamborg’s B5 1:10 medium amended with chitosan at different concentrations (0, 0.1, 1 and 2 mg·mL^-1^). Plants were then placed in a growth chamber at 24°C, 60% RH with a 16:8 hours light: dark photoperiod for 1, 3 and 5 days of incubation. This experiment was performed two times independently with ten biological replicates per treatment and time. Root exudates collected in this experiment are these accumulated over the indicated times. At 1, 3 and 5 days of incubation, plantlets were removed from the boxes and the exudates were collected and filtered through Miracloth (Merck Milipore, Calbiochem). The exudates of five biological replicates (5 mL) were mixed to obtain 2 pools per treatment to reduce the variability and concentrate the samples, filtered by 0.22 µm pore (Q-MAX) and frozen and stored at −20°C until use. To obtain the excitation emission matrix (EEM) fluorescence spectra we analysed two mL of each sample (pools of root exudates per treatment and time) with a spectrofluorometer (Jasco Model FP-6500)^26^. The EEM fluorescence spectra of the exudates were analysed by PARAFAC (Parallel Factor Analysis) with MATLAB (The MathWorks, Natick, MA)^97^. PARAFAC Model of Chemical Components were calculated for each treatment and time. The statistical analyses of the data obtained was performed using GraphPad Prism (8.1.2 version). The normality of the samples was checked using the Shapiro-Wilk Normality test and Levene’s Test to check the homogeneity of variances. In the case of not adjusting to a normal distribution, the Kruskal-Wallis non-parametric test was carried out, and if differences were found, the Wilcox test was done to analyse them. For those samples that adjusted to a normal distribution, ANOVA was performed, using a significance level of α = 0.05 to check if there were significant differences between treatments and, where appropriate, to identify them. Subsequently, an a posteriori test, TukeyHD test, was performed with two-by-two comparisons between treatments to observe between which groups the differences were, if any.

### Quantification of gene expression by RT-qPCR

Root material was collected from trays where plants were treated with chitosan (0 and 1 mg·mL^-1^) for 1 or 3 days. Roots from 6 plants per genotype, treatment and time point were snap frozen in liquid nitrogen and stored at −80lll°C. Total RNA was isolated^98^ while removal of gDNA traces and RNA quantity/quality checks were carried out^68^. For each DNA-free RNA sample 1 μg was reversed-transcribed to cDNA by using an oligo(dT)18 primer and the RevertAid H Minus First Strand cDNA Synthesis kit (Fermentas, St-Leon Rot, Germany) according to the manufacturer’s instructions.

Forty-one genes were selected to examine their root expression pattern and quantified in the trays system (Table S4). Out of the 41 genes selected in total, 8 genes are related to salicylic acid (SA)-defence, 5 genes related to jasmonic acid (JA)-defence, 6 genes associated with auxins biosynthesis, regulation, or transport, 4 genes involved in ethylene and ABA biosynthesis or regulation, 5 genes to reactive oxygen species (ROS) detoxification, 5 genes related to hypoxic stress, 4 genes to root growth and development and other 4 genes to parasitism and endophytism (Table S4). Primer sequences for 13 out of the 41 selected genes^80^, while for the remaining 28 genes the design and optimization of RT-qPCR primers was carried out as detailed in the above-mentioned study, to detect expression of all paralogs in each gene family. Primer combinations were custom ordered from commercial suppliers (Integrated DNA Technology, Coralville, IA, United States) and tested at two concentrations (100 and 150 nM) by gradient PCR with gDNA. Amplicon sizes were checked by 2% agarose gel electrophoresis and ethidium bromide staining.

RT-qPCR was carried out in 96-well plates with three technical replicates per sample using a StepOnePlusTM Real-Time PCR System (Applied Biosystems, USA). RT-qPCR design, calculations and statistics used followed the MIQE guidelines^99^. The master mix containing 1lll×lllABsoluteTM QPCR SYBR® Green Mix (Thermo Scientific, Epsom, UK), 100 nM of each forward and reverse primer (Supplementary Table S4) and 125lllng λ-DNA (Roche Diagnostics, Vilvoorde, Belgium) was mixed with 2lllμL of a 12x diluted sample cDNA, control gDNA or water. λ-DNA was added as carrier DNA to minimise absorption and Poisson effects. The following amplification program was used: 95lll°C for 15lllmin, 40 cycles of 95lll°C for 15llls, 60lll°C for 20llls and 72lll°C for 15llls. The melting curves of real-time qPCR products were assessed from 60 to 95°C to verify the specificity of the reactions under the following conditions: initial denaturation for 15 s at 95°C, cooling to 60°C for 1 min, and melting from 60 to 95°C with 0.3°C transition rate every 10 s. To calculate gene-specific amplification efficiencies (E), correlation coefficients (R^2^) and linear equations, standard curves were generated for each gene using five serial fourfold dilutions of sample-pooled cDNA and were included in each RT-qPCR run. The data obtained was normalized to the *M. acuminata Ribosomal protein L2* (*L2*) and *Elongation factor 1* α (*EF1*α) genes^68^ (Table S4). To determine gene expression levels, a baseline was set manually, and quantification cycles (Cq) values were converted into normalized relative quantities using qBASE + software (Biogazelle; Zwijnarde, Belgium)^100^. To check normality of gene expression data, the Shapiro–Wilk test was applied using STATISTICA 7.0 software (StatSoft, Inc., Tulsa, OK, United States). Data were multiplied by a common factor (10^3^) and log_10_-transformed either to fit a normal distribution or to improve normality. Subsequently, factorial ANOVA and Fisher’s least significant difference mean comparison (*p*< 0.05) were applied using STATISTICA 7.0 to compare chitosan treatment *vs.* control conditions in both analysed time-points (1 and 3 days).

### Preventive experiments to analyse root colonization of *Fusarium oxysporum* f. sp. cubense Tropical Race 4

*Fusarium oxysporum* f. sp. cubense Tropical Race 4 (ref. CBS 102025) and the nematophagous fungus *Pochonia chlamydosporia* strain 123 (ref. CBS 102025) isolated from eggs from *Heterodera avenae* ^101^ were used in our experiments. For conidia extraction, 30 g of rice colonised with a given fungus were added to 40 mL of sterile distilled water (SDW), vortexed 1 min and the suspension was filtered through sterile Miracloth (Merck Milano). A 10^6^ conidia·ml^-1^ suspension was added to the Magenta Box. *P. chlamydosporia*, inoculation was carried aout at the start of the treatments, while *F. oxysporum* TR4 was inoculated on the fifth day using 0.05% water agar conidia suspension. Fungi were grown on sterile rice substrate for by adding 4 cylinders of mycelium from the edge of a 10/21-day-old colony, grown in corn meal agar (CMA), of the corresponding fungus and 10 mL of SDW.

The experiment consisted of a hydroponic system using Magenta Boxes where plantlets of *Musa acuminata* were left with different solutions. Ten plantlets per treatment were used, placing two in each Magenta Box. Treatment volume solution was 50 mL. Banana plantlets were kept in a culture chamber with a photoperiod of 16h: 8h light: darkness, 24°C and 65% relative humidity, for 6 days. The experiment was carried out twice. The plants were exposed to different treatments: Control treatment (Gamborg’s); Chitosan treatment (1 mg mL^-1^); Control solution and Pc123 10^6^ spores mL^-1^; Chitosan 1 mg mL^-1^ and Pc123 10^6^ spores mL^-1^; Control solution and FocTR4 10^6^ spores mL^-1^ in 0.05% water agar; Chitosan 1 mg mL^-1^ and FocTR4 10^6^ spores mL^-1^ in 0.05% water agar; Control solution and Pc123 10^6^ spores mL^-1^ and FocTR4 10^6^ spores mL^-1^ in 0.05% water agar; Chitosan 1 mg mL^-1^ and Pc123 10^6^ spores mL^-1^ and FocTR4 10^6^ spores mL^-1^ in 0.05% aqueous agar. The plantlets were left growing 5 days with the different treatments without FocTR4, and then, at the sixth day the conidial solution of FocTR4 was added to the specific treatments, hence they are preventive treatments. After 24 hours, the plantlets were recovered for root colonization evaluation and DNA extraction.

Twenty-four hours after FocTR4 inoculation, roots were used for total and endophytic colonization assessments. From each treatment, three roots were used for evaluating total colonization, other three for endophytic colonization and the last four root samples were preserved frozen in liquid nitrogen and kept at −20°C until their use for DNA extraction. For total colonization, roots were rinsed in SDW three times for 1 min each. After that, they were blotted dry on sterile filter paper. Ten 10 mm fragments were collected randomly, axenically plated on CMA medium and incubated at 24°C to observe fungal growth. For endophytic colonization, roots were surface sterilised with 1% sodium hypochlorite for 1 min prior to the SDW rinse. They were then processed as for the total colonization assesments. Roots were evaluated for five consecutive days, scoring the number of colonized fragments each day, starting the observation period 24 hours after incubation. This served to evaluate the percentage of FocTR4 colonized fragments between the different treatments.

Banana roots and fungal mycelia, grown on liquid medium, were crushed to a fine powder (100 mg). DNA was extracted using DNeasy Plant Mini Kit (QIAGEN). DNA extraction of roots with chitosan treatments was carried out using the Acetyl-trimethyl-ammonium bromide (CTAB buffer: Tris 100 Mm, pH = 8.4; NaCl2 1.4 M; EDTA 25 Mm, pH = 7.5 and CTAB 2%+PVP 2%) method of O’Donnell^102^. Two quantitative PCRs (qPCR) were performed in triplicate using a 96-well plate, for FocTR4 and PC123 DNA quantification in banana roots with the different preventive treatments, in a Thermal Cycling StepOnePlus (Applied Biosystems). Primers used, specifics for each target, were: SIX1α (gene SIX1 for FocTR4 detection, F: CCCTCTCAATCCTTGGGTTT; R: TAGTGTCATTCCACGGCAAA, 153bp; PCR reaction: 95°C, 5 min; 35 cycles at 94°C, 30 sg, 58°C, 30 sg, 72°C, 30 sg^103^ and CSN1 (gene CSN1 for Pc123 detection, F: TCTTGCTGCTGTTACCTTGG; R: AGGAAGATGGCATTGGGAA, 222bp; PCR reaction: 95°C, 5 min; 35 cycles at 95°C, 30 sg, 30°C, 30 sg, 72°C, 30 sg; 95°C, 3 min^104^). For PCR reactions, the volumes of each reagent used were as follows: FastStart Universal SYBR Green Master (Roche) 5.50 μL; 0.5 μL of each primer; Nuclease-free water 2.5 μL; Sample (DNA) 1 μL. In addition to the quantitative reactions, two calibration curves were performed with DNA from each fungus, using serial dilutions 1:10 (100 ng·μl^-1^-0.001 ng·μl^-1^). Standard curves were plotted using values of the average cycle threshold (Ct values) against the logarithm of the DNA concentration (dilution) and, the amplification was calculated from the slope of this graph^105,106^. Prior to the quantitative PCRs, conventional PCRs were performed to evaluate the specificity of the primers, also EF1α (gene EF1α for *M. acuminata* detection, F: CCCACCGGTGCTAAGATCAC; R: AGGTGCCGATCAAACTGTCG, 128bp; PCR reaction: 95°C, 5 min; 40 cycles at 95°C, 15 sg, 60°C, 20 sg, 72°C, 15 sg^68^, was used to discriminate banana DNA too.

Root colonization results were analysed carrying out a 1-factor analysis of variance (ANOVA) to observe whether there were statistically significant differences between treatments using a significance level of α=0.05. Before performing the analysis, the normality and homoscedasticity of the data were checked by Shapiro-Wilk and Levene’s tests, respectively. Subsequently, an a posteriori test, Tukey HD test, was performed with two-by-two comparisons between treatments to observe between which groups the differences were, if any. The numerical data used to perform the analysis were those corresponding to the fifth day of incubation.

### Chitosan application by irrigation on *M. accuminata* plants

Banana plantlets were placed individually in 200 mL expanded polystyrene cups with sterilized (2 cycles at 121°C, 90 min, 1 atm) Terraplant II Sphagnum peat (Compo Expert) as plant substrate. Plants were grown for 30 days in a growth chamber at 24°C, 60% RH with a 16:8 hours light: dark photoperiod. Plants were treated differently by irrigating them with a sterilized solution of Gamborg’s B5 basal mixture 1:10 mixed with chitosan at different concentrations (0, 0.1, 1 and 2 mg·mL^-1^) every 2-3 days to field capacity. Plants were initially watered with 45 mL of the treatments’ solutions. Thirty days after planting plants were harvested for the measurement of developmental parameters (shoot and root length and weight and number of leaves). This experiment was repeated three times with 14 plantlets per treatment in each replicate. For the greenhouse experiment, firstly, we put the plants 30 days in a culture chamber and then passed them to a greenhouse for 2.5 months irrigating them every two weeks with 500 mL of specific treatments: watering Gamborg’s B5 basal mixture 1:10 and irrigate with Gamborg’s amended with chitosan at 1 mg·mL^-1^. Between treatments plants were drip irrigated. At the end of the experiment, we measured growth indicators as in the experiment explained before. Nineteen plants were used per treatment. The statistical study of the data obtained was performed using GraphPad Prism (8.1.2 version). The normality of the samples was checked using the Shapiro-Wilk Normality test and Levene’s Test to check the homogeneity of variances. In the case of not adjusting to a normal distribution, the Kruskal-Wallis non-parametric test was carried out, and if differences were found, the Wilcox test was done to analyse them. For those samples that adjusted to a normal distribution, ANOVA was performed, using a significance level of α = 0.05^92^ to check if there were significant differences between treatments and, where appropriate, to identify them. Subsequently, an a posteriori test, Tukey HD test, was performed with two-by-two comparisons between treatments to observe between which groups the differences were, if any.

## Acknowledgements

Authors would like to specially thank Ms Rocio Tirado-Conejo and Ms Maria Garcia-Mula (University of Alicante) for their technical assistance on FocTR4 related experiments, Ms Raquel Lopez-Nuñez (University of Alicante) on the banana plant growth experiments and Dr. Marta Suarez-Fernandez (Regional Service for Agrofood Research and Development (SERIDA), Spain) for her help on technical issues. Thanks also to Prof. Frutos Marhuenda (University of Alicante) for help on VOC data.

## Author contributions

Conceptualization and planification of the studies: F L-M, LV L-LL, A L-S, J Z-F, D GS, B G, JA LJ. Performance of the experiments and data collection: Root electrophysiology: A L-S, A H, A T, B G; Membrane lipid composition of fatty acids: F L-M, D GS, JA LJ; Analysis of VOCs: A L-S, N FLG; UHPLC hormone analysis and EEM fluorescence analysis: A L-S; Quantification of gene expression by RT-qPCR: J Z-F; Analysis of FocTR4 root colonization: F L-M, CM MG; Chitosan irrigation on banana plants: A L-S. Statistical analysis, data curation and analysis: A L-S, J Z-F, N FLG, CM MG, A H, A T, D GS, B G. Graphical visualization of the results: A L-S, J Z-F, B G, N FLG, CM MG. Results interpretation: F L-M, LV L-LL, A L-S, J Z-F, N FLG, CM MG, A H, A T, D GS, B G, JA LJ. Original draft writing and preparation: F L-M, LV L-LL, A L-S, J Z-F, N FLG, CM MG D GS. Draft editing and reviewing: F L-M, LV L-LL, A L-S, J Z-F, B G, JA LJ. Project administration: F L-M, LV L-LL. Funding acquisition: F L-M, LV L-LL. All authors have read and approved the manuscript.

## Additional information

This research was funded by PID2020-119734RB-I00 Project from the Spanish Ministry of Science and Innovation and by European Project H2020 MUSA no. 727624.

## Competing interests

The authors declare no conflict of interest.

## Notes

### Competing Interest Statement

The authors have declared no competing interest.

## References

1. Lescot T. (2011). La diversité génétique des bananiers, FruiTrop 89, 58–62.

2. Ferri, D. V., Munhoz, C. F., Neves, P. M. O., Ferracin, L. M., Sartori, D., Vieira, M. L. C., & Fungaro, M. H. P. (2012). Genetic variability of *Beauveria bassiana* and a DNA marker for environmental monitoring of a highly virulent isolate against *Cosmopolites sordidus*. Indian Journal of Microbiology, 52(4), 569–574.

3. Simmonds N.W. (1962) The Evolution of the Bananas. Longmans, Green and Co. Ltd., London, pp. 170.

4. Nelson S.C., Ploetz R.C., Kepler A.K. (2006). Musa species (banana and plantain). Species Profiles for Pacific Island agroforestry, 15.

5. Ostmark, H. E. (1974). Economic insect pests of bananas. Annual Review of Entomology, 19(1), 161–176.

6. Dubois, T., Gold, C. S., Coyne, D., Paparu, P., Mukwaba, E., Athman, S., Kapinduand, E. & Adipala, E. (2004). Merging biotechnology with biological control: Banana *Musa* tissue culture plants enhanced by endophytic fungi. Uganda Journal of Agricultural Sciences. 9(1), 445–451.

7. Waweru B., Turoop L., Kahangi E., Coyne D., Dubois T. (2014). Non-pathogenic *Fusarium oxysporum* endophytes provide field control of nematodes, improving yield of banana (*Musa* sp.). Biological Control 74, 82–88.

8. MAPA, Ministerio de Agricultura y Pesca, Alimentación y Medio Ambiente, MAPA (2021b) https://www.mapa.gob.es/es/agricultura/temas/sanidad-vegetal/productos-fitosanitarios/registro/productos/forexi.asp?e=0&plagEfecto=565 (Last checked 21dic2021).

9. Kaku, H., Nishizawa, Y., Ishii-Minami, N., Akimoto-Tomiyama, C., Dohmae, N., Takio, K., & Shibuya, N. (2006). Plant cells recognize chitin fragments for defense signaling through a plasma membrane receptor. Proceedings of the National Academy of Sciences 103(29), 11086–11091.

10. Dutta, P. K., Dutta, J., & Tripathi, V. S. (2004). Chitin and chitosan: Chemistry, properties, and applications. Journal of Scientific and Industry Research, 63, 20–31.

11. Lopez-Moya, F., Colom-Valiente, M.F., Martinez-Peinado, P., Martinez-Lopez, J.E., Puelles, E., Sempere-Ortells, J.M. & Lopez-Llorca, L.V. (2015). Carbon and nitrogen limitation increase chitosan antifungal activity in *Neurospora crassa* and fungal human pathogens. Fungal Biology 119 (2-3), 154–169.

12. Lopez-Moya, F., Escudero, N., Zavala-Gonzalez, E.A., Esteve-Bruna, D., Blazquez, M.A., Alabadi, D. & Lopez-Llorca, L.V. (2017). Induction of auxin biosynthesis and WOX5 repression mediate changes in root development in Arabidopsis exposed to chitosan. Scientific Reports 7, 16813.

13. Kumar, G., Bristow, J., Smith, P. & Payne, G. (2000). Enzymatic gelation of the natural polymer chitosan. Polymer 41(6), 2157–2168.

14. Palma-Guerrero, J., Jansson, H.-B., Salinas, J., & Lopez-Llorca, L. V. (2008). Effect of chitosan on hyphal growth and spore germination of plant pathogenic and biocontrol fungi. Journal of Applied Microbiology, 104(2), 541–553. 10.1111/j.1365-2672.2007.03567.x

15. Li, B., X. Wang, R. Chen, W. Huangfu & Xie, G.L. (2008). Antibacterial activity of chitosan solution against *Xanthomonas* pathogenic bacteria isolated from Euphorbia pulcherrima. Carbohydrate Polymers 72, 287–292.

16. Ferrante, P., & Scortichini, M. (2010). Molecular and phenotypic features of *Pseudomonas syringae* pv. actinidiae isolated during recent epidemics of bacterial canker on yellow kiwifruit (*Actinidia chinensis*) in central Italy. Plant Pathology 59, 954–962.

17. Benhamou, N., & Thériault, G. (1992). Treatment with chitosan enhances resistance of tomato plants to the crown and root rot pathogen *Fusarium oxysporum* f. sp. radicis-lycopersici. Physiological and Molecular Plant Pathology, 41(1), 33–52. 10.1016/0885-5765(92)90047-Y

18. Lafontaine P. & Benhamou N. (1996). Chitosan treatment: An emerging strategy for enhancing resistance of greenhouse tomato plants to infection by *Fusarium oxysporum* f.sp. radicis-lycopersici. Biocontrol Science and Technology 6, 111–124.

19. Trotel-Aziz P., Couderchet M., Vernet G. and Aziz A. (2006). Chitosan stimulates defense reactions in grapevine leaves and inhibits development of *Botrytis cinerea*. European Journal of Plant Pathology 114: 405–413.

20. Hassan O. and Chang T., (2017). Chitosan for Eco-friendly Control of Plant Disease. Asian Journal of Plant Pathology, 11: 53–70.

21. Lopez-Moya, F., Suarez-Fernandez, M., Lopez-Llorca, L.V. (2019). Molecular Mechanisms of Chitosan Interactions with Fungi and Plants. International Journal of Molecular Sciences 20(2):332

22. Chittenden, C., & Singh, T. (2009). In vitro evaluation of combination of *Trichoderma harzianum* and chitosan for the control of sapstain fungi. Biological Control, 50(3), 262–266.

23. López-Mondéjar, R., Blaya, J., Obiol, M., Ros, M., & Pascual, J. A. (2012). Evaluation of the effect of chitin-rich residues on the chitinolytic activity of *Trichoderma harzianum*: in vitro and greenhouse nursery experiments. Pesticide Biochemistry and Physiology 103(1), 1–8.

24. Melida, H., Sopeña-Torres, S., Bacete, L., Garrido-Arandia, M., Jordá, L., Lopez, G., Muñoz-Barrios, A., Pacios, L. F. & Molina, A. (2018). Non-branched β-1, 3-glucan oligosaccharides trigger immune responses in Arabidopsis. The Plant Journal, 93(1), 34–49.

25. El Hadrami, A., Adam, L.R., El Hadrami, I. and Daayf, F. (2010). Chitosan in Plant Protection, Marine Drugs, 8 (4), 968–987.

26. Suarez-Fernandez, M., Marhuenda-Egea, F. C., Lopez-Moya, F., Arnao, M. B., Cabrera-Escribano, F., Nueda, M. J., Lopez-Llorca, L. V. (2020). Chitosan induces plant hormones and defenses in tomato root exudates. Frontiers in Plant Science, 11, 572087.

27. Maruri-López, I., Aviles-Baltazar, N. Y., Buchala, A., & Serrano, M. (2019). Intra and extracellular journey of the phytohormone salicylic acid. Frontiers in Plant Science, 423, 1–11.

28. Snoeren, T. A., Mumm, R., Poelman, E. H., Yang, Y., Pichersky, E., & Dicke, M. (2010). The herbivore-induced plant volatile methyl salicylate negatively affects attraction of the parasitoid *Diadegma semiclausum*. Journal of Chemical Ecology, 36(5), 479–489.

29. Lefevere, H., Bauters, L., & Gheysen, G. (2020). Salicylic acid biosynthesis in plants. Frontiers in Plant Science, 11, 338.

30. Shulaev, V., Silverman, P., & Raskin, I. (1997). Airborne signalling by methyl salicylate in plant pathogen resistance. Nature, 385(6618), 718–721.

31. Chen, F., D’Auria, J. C., Tholl, D., Ross, J. R., Gershenzon, J., Noel, J. P., & Pichersky, E. (2003). An *Arabidopsis thaliana* gene for methyl salicylate biosynthesis, identified by a biochemical genomics approach, has a role in defense. The Plant Journal, 36(5), 577–588.

32. Attaran, E., Zeier, T. E., Griebel, T., & Zeier, J. (2009). Methyl salicylate production and jasmonate signaling are not essential for systemic acquired resistance in Arabidopsis. The Plant Cell, 21(3), 954–971.

33. Heil, M., Walters, D., Newton, A., & Lyon, G. (2014). Trade-offs associated with induced resistance. Induced Resistance for Plant Defense: A Sustainable Approach to Crop Protection, eds DR Walters, AC Newton, and GD Lyon (West Sussex: John Wiley & Sons, Ltd.), 171-185.

34. Masimbula, R., Oki, K., Takahashi, K., & Matsuura, H. (2020). Metabolism of airborne methyl salicylate in adjacent plants. Bioscience, Biotechnology, and Biochemistry, 84(9), 1780–1787.

35. Gao, Q. M., Zhu, S., Kachroo, P., & Kachroo, A. (2015). Signal regulators of systemic acquired resistance. Frontiers in Plant Science 6, 228.

36. Suwanchaikasem, P., Nie, S., Selby-Pham, J., Walker, R., Boughton, B.A., Idnurm, A., (2023) Hormonal and proteomic analyses of southern blight disease caused by and root chitosan priming on in an in vitro hydroponic system, Plant Direct, 7, 9.

37. Cassim, A. M., Gouguet, P., Gronnier, J., Laurent, N., Germain, V., Grison, M., & Mongrand, S. (2019). Plant lipids: Key players of plasma membrane organization and function. Progress in Lipid Research, 73, 1–27.

38. Brilli, F., Ruuskanen, T. M., Schnitzhofer, R., Müller, M., Breitenlechner, M., Bittner, V., & Hansel, A. (2011). Detection of plant volatiles after leaf wounding and darkening by proton transfer reaction “time-of-flight” mass spectrometry (PTR-TOF). PLoS One, 6(5), e20419.

39. Dudareva, N., Negre, F., Nagegowda, D. A., & Orlova, I. (2006). Plant volatiles: recent advances and future perspectives. Critical Reviews in Plant Sciences, 25(5), 417–440.

40. Portillo-Estrada, M., & Niinemets, Ü. (2018). Massive release of volatile organic compounds due to leaf midrib wounding in *Populus tremula*. Plant Ecology, 219(9), 1021–1028.

41. Scala, A., Mirabella, R., Mugo, C., Matsui, K., Haring, M. A., & Schuurink, R. C. (2013). E-2-hexenal promotes susceptibility to *Pseudomonas syringae* by activating jasmonic acid pathways in Arabidopsis. Frontiers in Plant Science, 4, 74.

42. Keinath, N. F., Kierszniowska, S., Lorek, J., Bourdais, G., Kessler, S. A., Shimosato-Asano, H., Grossniklaus, U., Schulze, W. X., Robatzek, S., & Panstruga, R. (2010). PAMP (pathogen-associated molecular pattern)-induced changes in plasma membrane compartmentalization reveal novel components of plant immunity. The Journal of Biological Chemistry, 285(50), 39140–39149.

43. Walley, J. W., Kliebenstein, D. J., Bostock, R. M., & Dehesh, K. (2013). Fatty acids and early detection of pathogens. Current opinion in plant biology, 16(4), 520–526.

44. Živković, S., Skorić, M., Ristić, M., Filipović, B., Milutinović, M., Perišić, M., & Puač, N. (2021). Rehydration Process in Rustyback Fern (*Asplenium ceterach* L.): Profiling of Volatile Organic Compounds. Biology, 10(7), 574.

45. Reszczyńska, E., & Hanaka, A. (2020). Lipids composition in plant membranes. Cell Biochemistry and Biophysics, 78(4), 401–414.

46. Raffaele, S., Leger, A., & Roby, D. (2009). Very long chain fatty acid and lipid signaling in the response of plants to pathogens. Plant Signaling & Behavior, 4(2), 94–99.

47. Martínez, G.; Olivares, B.O.; Rey, J.C.; Rojas, J.; Cardenas, J.; Muentes, C.; Dawson, C. (2023). The Advance of *Fusarium* Wilt Tropical Race 4 in Musaceae of Latin America and the Caribbean: Current Situation. Pathogens 12(2), 277.

48. Doares, S. H., Syrovets, T., Weiler, E. W., & Ryan, C. A. (1995). Oligogalacturonides and chitosan activate plant defensive genes through the octadecanoid pathway. Proceedings of the National Academy of Sciences, 92(10), 4095–4098.

49. Chandra, S., Chakraborty, N., Dasgupta, A., Sarkar, J., Panda, K., & Acharya, K. (2015). Chitosan nanoparticles: a positive modulator of innate immune responses in plants. Scientific Reports, 5(1), 1–14.

50. Lopez-Moya, F., & Lopez-Llorca, L. V. (2016). Omics for investigating chitosan as an antifungal and gene modulator. Journal of Fungi 2(1), 11.

51. Escudero, N., Ferreira, S. R., Lopez-Moya, F., Naranjo-Ortiz, M. A., Marin-Ortiz, A. I., Thornton, C. R., & Lopez-Llorca, L. V. (2016). Chitosan enhances parasitism of *Meloidogyne javanica* eggs by the nematophagous fungus *Pochonia chlamydosporia*. Fungal Biology, 120(4), 572–585.

52. Escudero, N., Lopez-Moya, F., Ghahremani, Z., Zavala-Gonzalez, E. A., Alaguero-Cordovilla, A., Ros-Ibañez, C., & Lopez-Llorca, L. V. (2017). Chitosan increases tomato root colonization by *Pochonia chlamydosporia* and their combination reduces root-knot nematode damage. Frontiers in Plant Science, 8, 1415.

53. Palma-Guerrero, J., Lopez-Jimenez, J.A., Perez-Berna, A.J., Huang, I., Jansson, H., Salinas, J., Villalain, J., Read, N.D. and Lopez-Llorca, L.V. (2010). Membrane fluidity determines sensitivity of filamentous fungi to chitosan, Molecular Microbiology, vol. 75, no. 4, pp. 1021–1032.

54. Meisrimler, C. N., Planchon, S., Renaut, J., Sergeant, K., & Lüthje, S. (2011). Alteration of plasma membrane-bound redox systems of iron deficient pea roots by chitosan. Journal of Proteomics, 74(8), 1437–1449.

55. Jaime, M. D., Lopez-Llorca, L. V., Conesa, A., Lee, A. Y., Proctor, M., Heisler, L. E., & Nislow, C. (2012). Identification of yeast genes that confer resistance to chitosan oligosaccharide (COS) using chemogenomics. BMC Genomics 13(1), 1–26.

56. He, W., Guo, X., Xiao, L., & Feng, M. (2009). Study on the mechanisms of chitosan and its derivatives used as transdermal penetration enhancers. International Journal of Pharmaceutics 382(1-2), 234–243.

57. Pandey, G.K. (2017). Mechanism of Plant Hormone Signaling Under Stress. Hoboken, NJ: John Wiley & Sons.

58. Wasternack C. (2007). Jasmonates: an update on biosynthesis, signal transduction and action in plant stress response, growth, and development. Annals of Botany 100(4):681–97.

59. Wasternack, C., & Hause, B. (2013). Jasmonates: biosynthesis, perception, signal transduction and action in plant stress response, growth, and development. An update to the 2007 review in Annals of Botany. Annals of botany, 111(6), 1021–1058.

60. Dave, A., & Graham, I. A. (2012). Oxylipin signaling: a distinct role for the jasmonic acid precursor cis-(+)-12-oxo-phytodienoic acid (cis-OPDA). Frontiers in Plant Science, 3, 42.

61. Noverr, M. C., Erb-Downward, J. R., & Huffnagle, G. B. (2003). Production of eicosanoids and other oxylipins by pathogenic eukaryotic microbes. Clinical microbiology reviews, 16(3), 517–533.

62. ul Hassan, M. N., Zainal, Z., & Ismail, I. (2015). Green leaf volatiles: biosynthesis, biological functions, and their applications in biotechnology. Plant Biotechnology Journal, 13(6), 727–739.

63. Spyropoulou, E. A., Dekker, H. L., Steemers, L., van Maarseveen, J. H., de Koster, C. G., Haring, M. A., Schuurink, R.C. and Allmann, S. (2017). Identification and characterization of (3 Z): (2 E)-hexenal isomerases from cucumber. Frontiers in Plant Science, 8, 1342.

64. Cui, H., Gobbato, E., Kracher, B., Qiu, J., Bautor, J., & Parker, J. E. (2017). A core function of EDS1 with PAD4 is to protect the salicylic acid defense sector in Arabidopsis immunity. New Phytologist, 213(4), 1802–1817.

65. Iriti, M., & Faoro, F. (2009). Chitosan as a MAMP, searching for a PRR. Plant Signaling & Behavior 4(1), 66–68.

66. Dar, T. A., Uddin, M., Khan, M. M. A., Hakeem, K. R., & Jaleel, H. (2015). Jasmonates counter plant stress: a review. Environmental and Experimental Botany, 115, 49–57.

67. Zhang, R. Q., Zhu, H. H., Zhao, H. Q., & Yao, Q. (2013). Arbuscular mycorrhizal fungal inoculation increases phenolic synthesis in clover roots via hydrogen peroxide, salicylic acid and nitric oxide signaling pathways. Journal of Plant Physiology, 170(1), 74–79.

68. Zorrilla-Fontanesi, Y., Rouard, M., Cenci, A., Kissel, E., Do, H., Dubois, E., & Carpentier, S. C. (2016). Differential root transcriptomics in a polyploid non-model crop: the importance of respiration during osmotic stress. Scientific Reports, 6(1), 1–15.

69. Buregyeya, H., Tumuhimbise, R., Matovu, M., Tumwesigye, K. S., Kubiriba, J., Nowankunda, K., Tushemereirwe, W. K., Karamura, D., Karamura, E., Kityo, R. M., & Rubaihayo, P. (2020). *Fusarium oxysporum* Race 1 resistance and quality traits variations in apple banana germplasm. Journal of Plant Breeding and Crop Science, 12(1), 16–24.

70. Mishina, T. E., & Zeier, J. (2006). The Arabidopsis flavin-dependent monooxygenase FMO1 is an essential component of biologically induced systemic acquired resistance. Plant Physiology, 141(4), 1666–1675.

71. Maruri-López, I., Aviles-Baltazar, N. Y., Buchala, A. & Serrano, M. (2019). Intra and extracellular journey of the phytohormone salicylic acid. Frontiers in Plant Science, 423

72. Wu, Y., Zhang, D., Chu, J. Y., Boyle, P., Wang, Y., Brindle, I. D., De Luca, V., & Després, C. (2012). The Arabidopsis NPR1 protein is a receptor for the plant defense hormone salicylic acid. Cell reports, 1(6), 639–647.

73. Pruitt, R. N., Locci, F., Wanke, F., Zhang, L., Saile, S. C., Joe, A., Karelina, D., Hua, C., Fröhlich, K., Wan, W. L., Hu, M., Rao, S., Stolze, S. C., Harzen, A., Gust, A. A., Harter, K., Joosten, M. H. A. J., Thomma, B. P. H. J., Zhou, J. M., Dangl, J. L., Nürnberger, T. (2021). The EDS1-PAD4-ADR1 node mediates Arabidopsis pattern-triggered immunity. Nature, 598(7881), 495–499.

74. Zhan, N.; Kuang, M.; He, W.; Deng, G.; Liu, S.; Li, C.; Roux, N.; Dita, M.; Yi, G.; Sheng, O. (2022). Evaluation of Resistance of Banana Genotypes with AAB Genome to Fusarium Wilt Tropical Race 4 in China. Journal of Fungi, 8, 1274.

75. Chen, A., Sun, J., Matthews, A., Armas-Egas, L., Chen, N., Hamill, S., Mintoff, S., Tran-Nguyen, L. T. T., Batley, J., & Aitken, E. A. B. (2019). Assessing Variations in Host Resistance to *Fusarium oxysporum* f sp. cubense Race 4 in *Musa* Species, With a Focus on the Subtropical Race 4. Frontiers in Microbiology, 10, 1062.

76. Dita, M., Barquero, M., Heck, D., Mizubuti, E. S. G., & Staver, C. P. (2018). *Fusarium* Wilt of Banana: Current Knowledge on Epidemiology and Research Needs Toward Sustainable Disease Management. Frontiers in Plant Science, 9, 1468.

77. Bubici, G., Kaushal, M., Prigigallo, M.I., Gómez-Lama Cabanás, C., & Mercado-Blanco, J. (2019). Biological Control Agents Against Fusarium Wilt of Banana. Frontiers in Microbiology 10, 616.

78. Wang, J., Cai, B., Li, K., Zhao, Y., Li, C., Liu, S., Xiang, D., Zhang, L., Xie, J., & Wang, W. (2022). Biological Control of *Fusarium oxysporum* f. sp. cubense Tropical Race 4 in Banana Plantlets Using Newly Isolated *Streptomyces* sp. WHL7 from Marine Soft Coral. Plant disease, 106(1), 254–259.

79. Li, Y., Jiang, S., Jiang, J., Gao, C., Qi, X., Zhang, L., Sun, S., Dai, Y., & Fan, X. (2022). Synchronized Efficacy and Mechanism of Alkaline Fertilizer and Biocontrol Fungi for *Fusarium oxysporum* f. sp. cubense Tropical Race 4. Journal of Fungi 8(3), 261.

80. Cabanás, C. G. L., Wentzien, N. M., Zorrilla-Fontanesi, Y., Valverde-Corredor, A., Fernández-González, A. J., Fernández-López, M., & Mercado-Blanco, J. (2022). Impacts of the Biocontrol Strain *Pseudomonas simiae* PICF7 on the Banana Holobiont: Alteration of Root Microbial Co-occurrence Networks and Effect on Host Defense Responses. Frontiers in Microbiology, 13.

81. Gómez-Lama Cabanás, C., Fernández-González, A. J., Cardoni, M., Valverde-Corredor, A., López-Cepero, J., Fernández-López, M., & Mercado-Blanco, J. (2021). The banana root endophytome: differences between mother plants and suckers and evaluation of selected bacteria to control *Fusarium oxysporum* f. sp. cubense. Journal of Fungi 7(3), 194.

82. Al-Hetar, M. Y., Zainal Abidin, M. A., Sariah, M., & Wong, M. Y. (2011). Antifungal activity of chitosan against *Fusarium oxysporum* f. sp. cubense. Journal of Applied Polymer Science, 120(4), 2434–2439.

83. Morán-Diez, E., Rubio, B., Domínguez, S., Hermosa, R., Monte, E., & Nicolás, C. (2012). Transcriptomic response of *Arabidopsis thaliana* after 24 h incubation with the biocontrol fungus *Trichoderma harzianum*. Journal of Plant Physiology, 169(6), 614–620.

84. Zavala-Gonzalez, E. A., Rodríguez-Cazorla, E., Escudero, N., Aranda-Martinez, A., Martínez-Laborda, A., Ramírez-Lepe, M., Vera, A., & Lopez-Llorca, L. V. (2017). *Arabidopsis thaliana* root colonization by the nematophagous fungus *Pochonia chlamydosporia* is modulated by jasmonate signaling and leads to accelerated flowering and improved yield. New Phytologist, 213(1), 351–364.

85. Mestre-Tomás, J.; Esgueva-Vilà, D.; Fuster-Alonso, A.; Lopez-Moya, F.; Lopez-Llorca, L.V. (2023) Chitosan Modulates Volatile Organic Compound Emission from the Biocontrol Fungus *Pochonia chlamydosporia*. Molecules, 28, 4053.

86. Chitarra, G. S., Abee, T., Rombouts, F. M., & Dijksterhuis, J. (2005). 1-Octen-3-ol inhibits conidia germination of *Penicillium paneum* despite of mild effects on membrane permeability, respiration, intracellular pH, and changes the protein composition. FEMS Microbiology Ecology, 54(1), 67–75.

87. Miyamoto, K., Murakami, T., Kakumyan, P., Keller, N. P., & Matsui, K. (2014). Formation of 1-octen-3-ol from *Aspergillus flavus* conidia is accelerated after disruption of cells independently of PPO oxygenases, and is not a main cause of inhibition of germination. Peer Journal, 2, e395.

88. Li, K., Xing, R., Liu, S., & Li, P. (2020). Chitin and chitosan fragments responsible for plant elicitor and growth stimulator. Journal of Agricultural and Food Chemistry 68(44), 12203–12211.

89. Chakraborty, M., Hasanuzzaman, M., Rahman, M., Khan, M. A. R., Bhowmik, P., Mahmud, N. U., Tanveer, M. & Islam, T. (2020). Mechanism of plant growth promotion and disease suppression by chitosan biopolymer. Agriculture, 10(12), 624.

90. van Wesemael, J., Hueber, Y., Kissel, E., Campos, N., Swennen, R., & Carpentier, S. (2018). Homeolog expression analysis in an allotriploid non-model crop via integration of transcriptomics and proteomics. Scientific Reports, 8(1), 1–11.

91. Gamborg, O. L., Miller, R., & Ojima, K. (1968). Nutrient requirements of suspension cultures of soybean root cells. Experimental Cell Research 50(1), 151–158.

92. Gunsé, B., Poschenrieder, C., Rankl, S., Schröeder, P., Rodrigo-Moreno, A., & Barceló, J. (2016). A highly versatile and easily configurable system for plant electrophysiology. Methods 3, 436–451.

93. Underwood, A. J. (1997). Experiments in ecology: their logical design and interpretation using analysis of variance. Journal of Environmental Quality, vol. 24, pp. 246.

94. Folch, J., Lees, M., & Sloane Stanley, G. H. (1957). A simple method for the isolation and purification of total lipids from animal tissues. Journal of Biological Chemistry 226(1), 497–509.

95. Christie, W.W., 2003. Lipid Analysis, third ed. The Oily Press, Bridgewater, UK, pp. 205–224.

96. Grafit, A., Muller, D., Kimchi, S., & Avissar, Y. Y. (2018). Development of a solid phase microextraction (SPME) Fiber protector and its application in flammable liquid residues analysis. Forensic Science International 292, 138–147.

97. Ohno, T., & Bro, R. (2006). Dissolved organic matter characterization using multiway spectral decomposition of fluorescence landscapes. Soil Science Society of America Journal, 70(6), 2028–2037.

98. Podevin, N., Krauss, A., Henry, I., Swennen, R., & Remy, S. (2012). Selection and validation of reference genes for quantitative RT-PCR expression studies of the non-model crop Musa. Molecular Breeding, 30(3), 1237–1252.

99. Bustin, S. A., Benes, V., Garson, J. A., Hellemans, J., Huggett, J., Kubista, M., Nolan, T., Pfaffl, M. W., Shipley, G. L., Vandesompele, J., & Wittwer, C. T. (2009). The MIQE Guidelines: Minimum Information for Publication of Quantitative Real-Time PCR Experiments. Clinical Chemistry, 55(4), 611–622.

100. Hellemans, J., Mortier, G., De Paepe, A., Speleman, F., & Vandesompele, J. (2007). Base relative quantification framework and software for management and automated analysis of real-time quantitative PCR data. Genome Biology 8(2), 1–14.

101. Olivares-Bernabeu, C.M.; López-Llorca, L.V. (2002). Fungal Egg-Parasites of Plant-Parasitic Nematodes from Spanish Soils. Rev. Iberoam. Micol. 19, 104–110.

102. O’Donnell, K.; Cigelnik, E.; Nirenberg, H.I. (1998). Molecular systematics and phylogeography of the *Gibberella fujikuroi* species complex. Mycologia, 90, 465– 493.

103. Maldonado-Bonilla, L. D., Calderón-Oropeza, M. A., Villarruel-Ordaz, J. L., & Sánchez-Espinosa, A. C. (2019). Identification of novel potential causal agents of *Fusarium* wilt of Musa sp. AAB in southern Mexico. Journal of Plant Pathology and Microbiology 10, e479.

104. Aranda-Martinez, A., Lenfant, N., Escudero, N., Zavala-Gonzalez, E. A., Henrissat, B., & Lopez-Llorca, L. V. (2016). CAZyme content of *Pochonia chlamydosporia* reflects that chitin and chitosan modification are involved in nematode parasitism: CAZome of *Pochonia chlamydosporia*. Environmental Microbiology, 18(11), 4200–4215.

105. Maciá-Vicente, J.G.; Jansson, H.-B.; Talbot, N.J.; Lopez-Llorca, L.V. (2009). Real-Time PCR Quantification and Live-Cell Imaging of Endophytic Colonization of Barley (*Hordeum vulgare*) Roots by *Fusarium equiseti* and *Pochonia chlamydosporia*. New Phytologist 182, 213–228.

106. Escudero, N., & Lopez-Llorca, L.V. (2012). Effects on Plant Growth and Root-Knot Nematode Infection of an Endophytic GFP Transformant of the Nematophagous Fungus *Pochonia chlamydosporia*. Symbiosis, 57, 33–42.

## References

1. Fernández-Falcón M. Borges, A. A. and Borges-Pérez, A. 2004. Response of Dwarf Cavendish banana plantlets to inoculation with races 1 and 4 of Fusarium oxysporum f. sp. cubense at differen Zn nutrition. Fruits 59: 319–323

2. Carlier J.. D. De Waele and J.V. Escalant. 2003. Global evaluation of Musa germplasm for resistance to Fusarium wilt. Mycosphaerella leaf spot diseases and nematodes. Performance evaluation (A nd C. Picq. eds). INIBAP Technical Guidelines 7. The International Network for the Improvement of Banana and Plantain. Montpellier. France.

3. Molina, A.B., et al. 2016. Resistance to Fusarium oxysporum f. sp. cubense tropical race 4 in African bananas. Acta Hortic. 1114

4. Ploetz, R.C. Haynes, J. L. and Vázuez, A. 1999. Responses of new banana accessions in South Florida to Panama disease. Crop Protection 18: 445–449

5. Tushemereirwe, W.K. Kangire, A. Kubiriba, J. and Nowakunda, K. 2000. Fusarium wilt resistant bananas considered appropriate replacements for cultivars susceptible to the disease in Uganda ournal of Agricultural Sciences 5:62–64

6. Zuo, C., et al.. 2018. Germplasm screening of Musa spp. for resistance to Fusarium oxysporum f. sp. cubense tropical race 4 (Foc TR4). European Journal of Plant Pathology 151:723–734

