## Supplementary Material for "Chitosan induces salicylic acid local and systemically in banana plants and reduces colonization by the pathogen *Fusarium oxysporum* f. sp. cubense TR4"

**Supplementary figures**

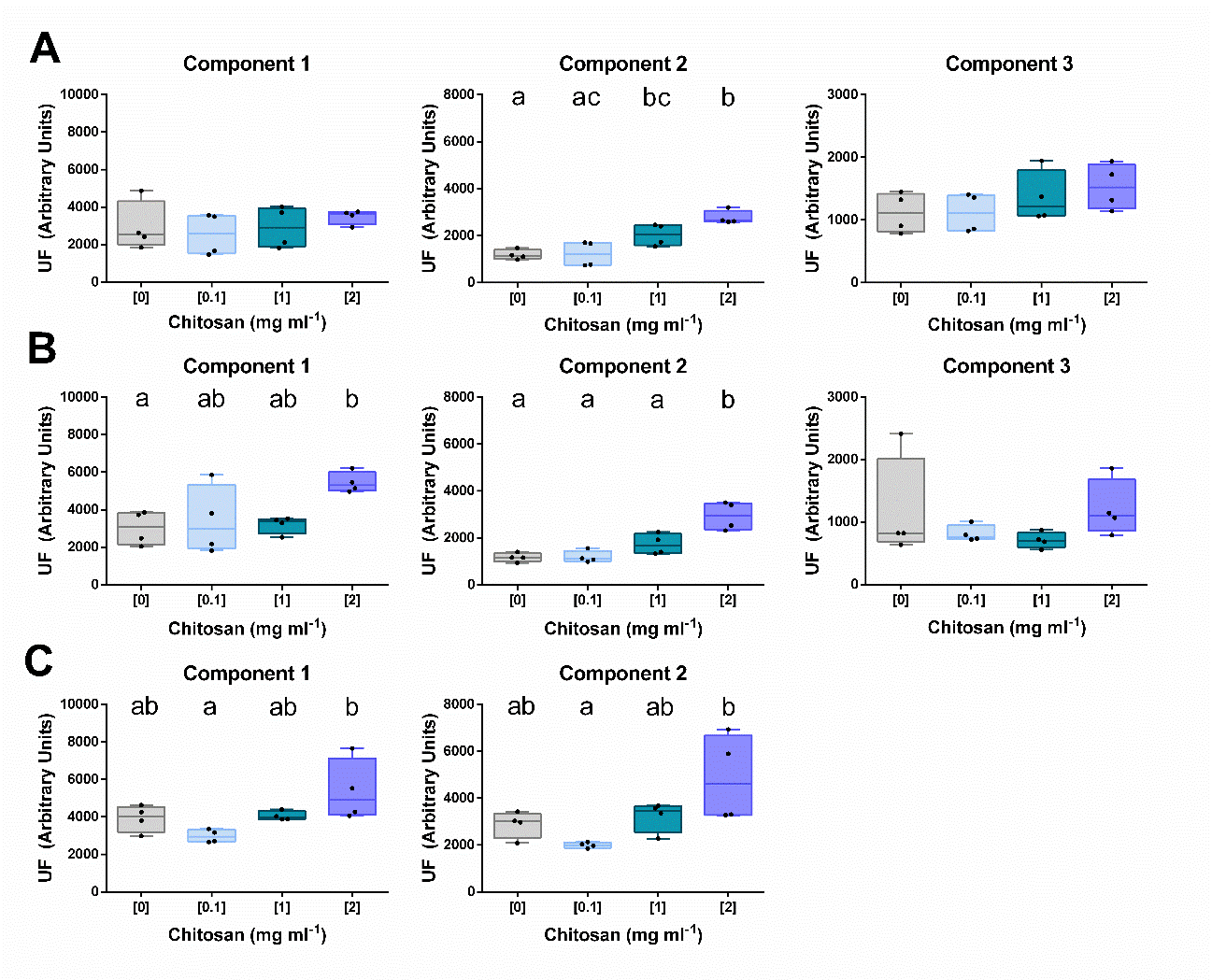

**Figure S1. Chitosan modifies EEM fluorescence of banana (cv. Petite Naine) root exudates.** The analysis showed 3 principal components (CORCONDIA >80%; Table S1). Components 1 and 2 were always the most induced (1. 3 and 5 days) after chitosan treatments (0. 0.1. 1 and 2 mg·ml^-1^). **A. 1-day-chitosan treatment.** Component 2 increases significantly its fluorescence in root exudates with higher doses of chitosan. **B. 3-days-chitosan treatment.** The fluorescence of components 1 and 2 increases which higher doses of chitosan. **C. 5-days-chitosan treatment.** Component 3 does not appear in the PARAFAC model. whereas components 1 and 2 follow the same tendency of the previous times. Two replicates. each one with two pools of five root exudates samples from different plants were analyzed. Different letters indicate significant differences based on comparison over treatments (ANOVA; Tukey: α = 0.05). Component 1: possible indole acetic acid; Component 2: possible salicylic acid derivatives and phenolics; Component 3: possible phenolics and aromatic amino acids and peptides (more data in Table S1).

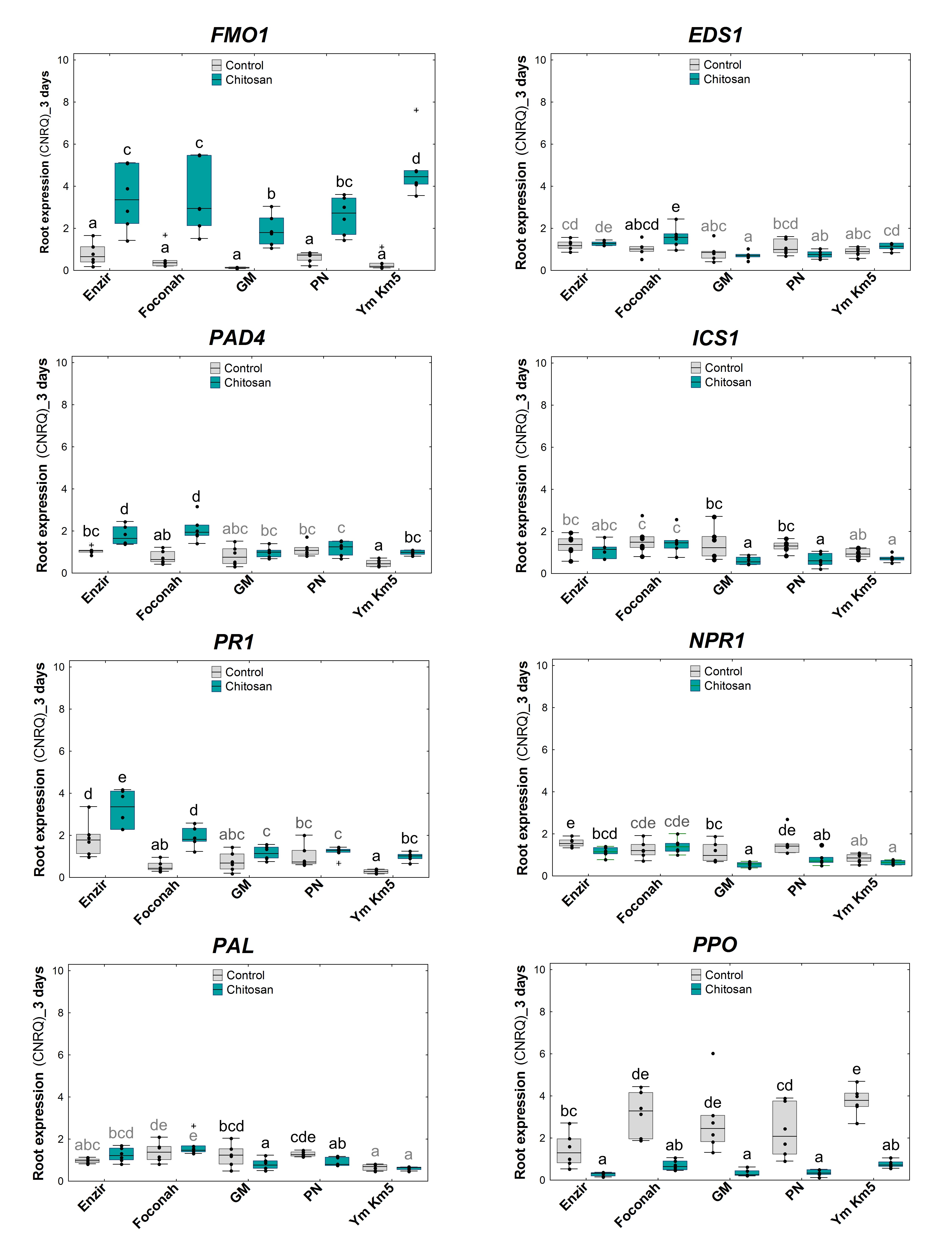

**Figure S2. Chitosan induces expression of genes involved in salicylic acid (SA) metabolism and SAR in five banana genotypes exposed 3 days to chitosan treatment.** Relative gene expression levels quantified in banana roots of diverse genotypes subjected to chitosan treatment in hydroponic tray system. **B).** CNRQ: Calibrated Normalized Relative Quantity. *Musa* genes *EF-1* and *ACT-1* were used as internal controls to normalize the expression data. Error bars = standard error of the mean (SEM). Different letters indicate significant differences based on comparison over treatments. genotypes and genotype x treatment (Two-way ANOVA and Tukey; α = 0.05). N control/treatment= 6/6. Gene abbreviations according to Table S3.

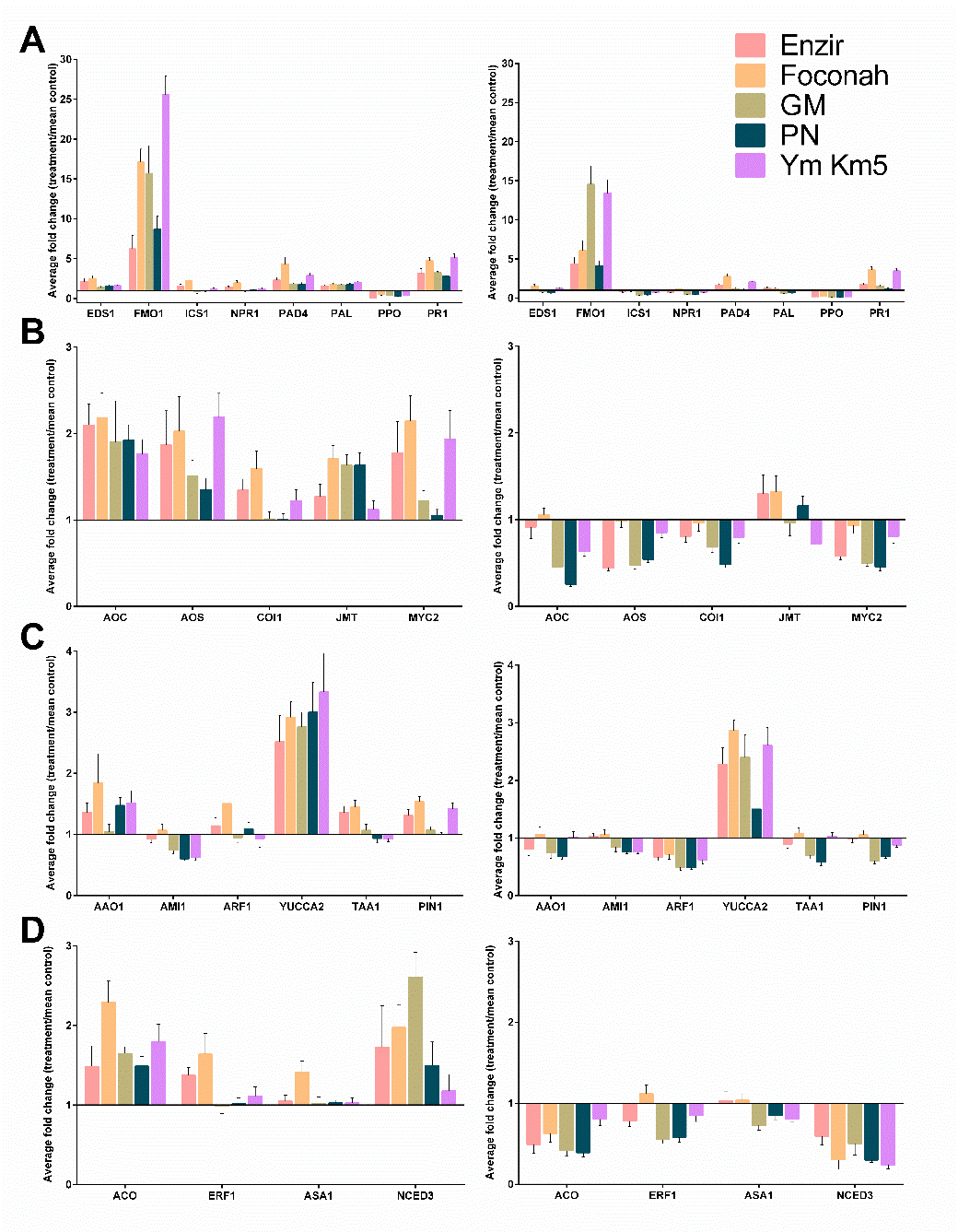

**Figure S3. Relative root expression (fold changes) of the 41 genes quantified in five banana genotypes subjected to chitosan treatment (1 mg·ml^-1^) in trays system.** Genes related **to A)** Salicylic acid defense; **B)** Jasmonic defense; **C**) Auxin biosynthesis. signaling or transport; and **D)** Ethylene or ABA biosynthesis. signaling or transport**.** *Musa* genes *EF-1* and *L2* were used as internal controls to normalize the expression. Enzir: Enzirabahima; GM: Gros Michel; PN: Petite Naine; Ym Km5: Yangambi Km5. Error bars: standard error of the means (SEM). N control/treatment=6/6. 1 day (left) and 3 days (right)

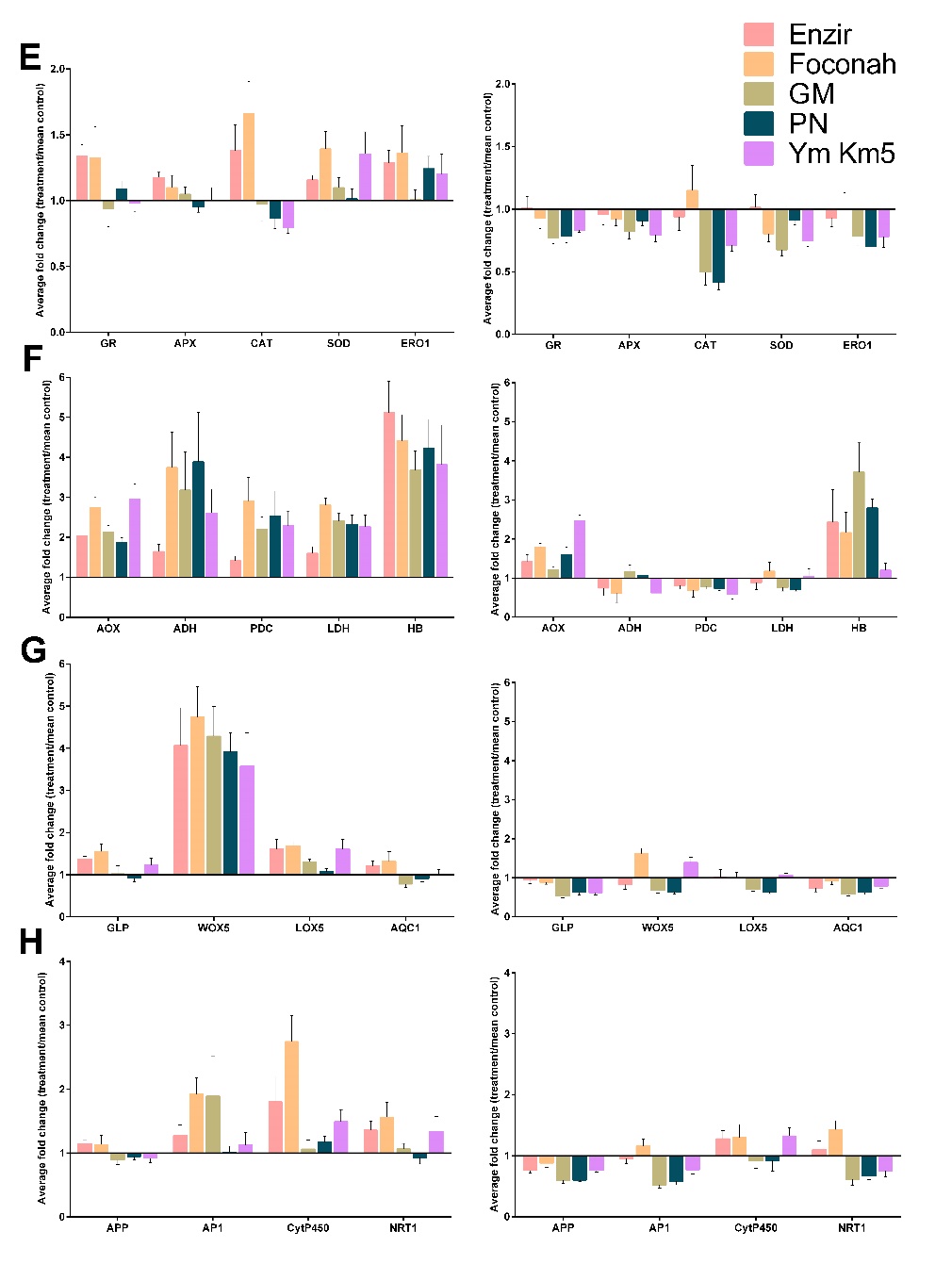

**Figure S3. Continued. Relative root expression (fold changes) of the 41 genes quantified by RT-qPCR in five different banana genotypes after chitosan (1 mg·mL^-1^) treatment in trays system. Genes related to E**) ROS detoxification; **F)** Hypoxia; **G)** Root growth and development; and **H)** Parasitism-endophytism. *Musa* genes *EF-1* and *L2* were used as internal controls to normalize the expression. Enzir: Enzirabahima; GM: Gros Michel; PN: Petite Naine; Ym Km5: Yangambi Km5. Error bars: standard error of the means (SEM). d: day(s) after treatment. N control/treatment=6/6.

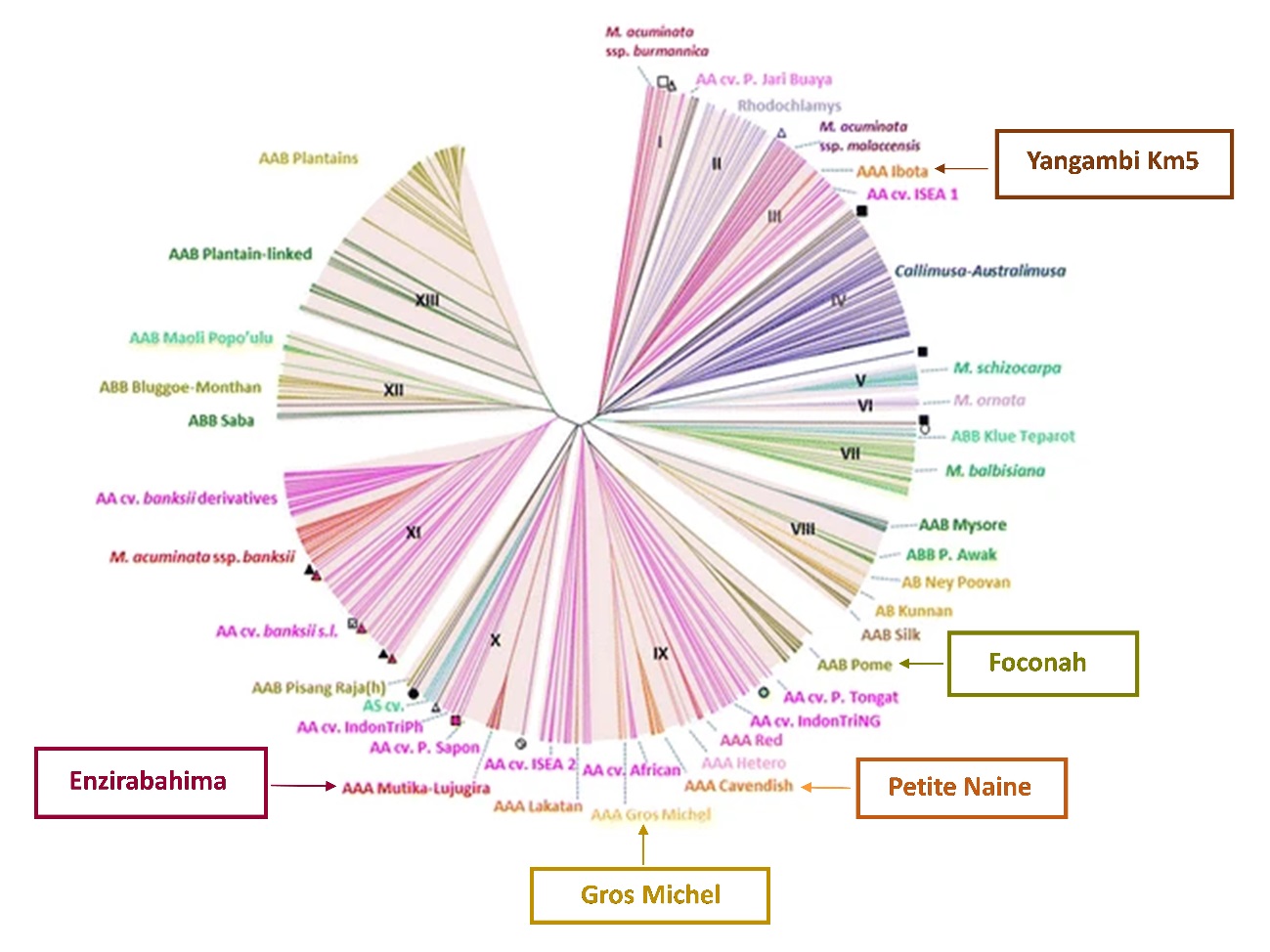

**Figure S4. Location on the *Musa* core subset dendrogram of the five banana genotypes selected for this study.** Adapted from Christelová. P.. De Langhe. E.. Hřibová. E. *et al*. (2017). Molecular and cytological characterization of the global *Musa* germplasm collection provides insights into the treasure of banana diversity. Biodivers. Conserv. 26. 801–824. <https://doi.org/10.1007/s10531-016-1273-9>

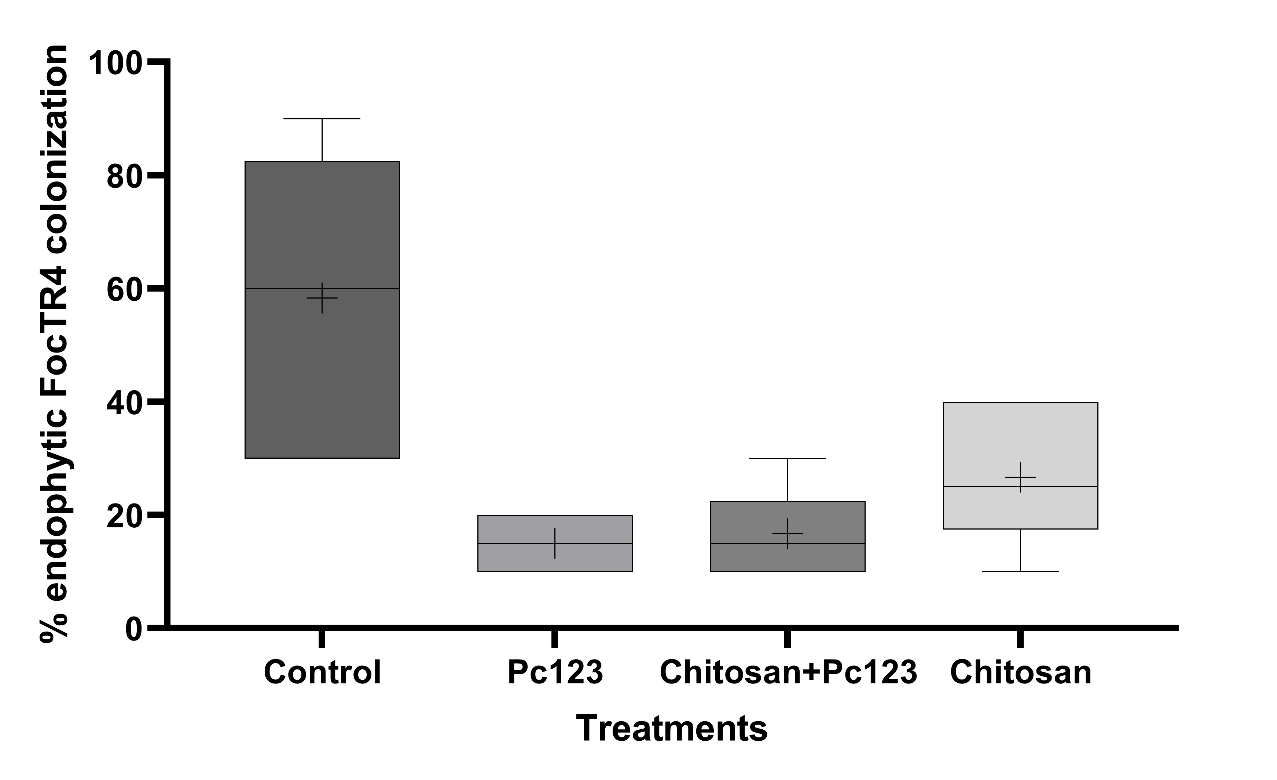

**Figure S5. Quantification of fungal colonization of banana (cv. Petit Naine) roots.** Quantification of *P. chlamydosporia* strain 123 and *Fusarium oxysporum* f. sp. *cubense* TR4 endophytic root colonization by culturing techniques.

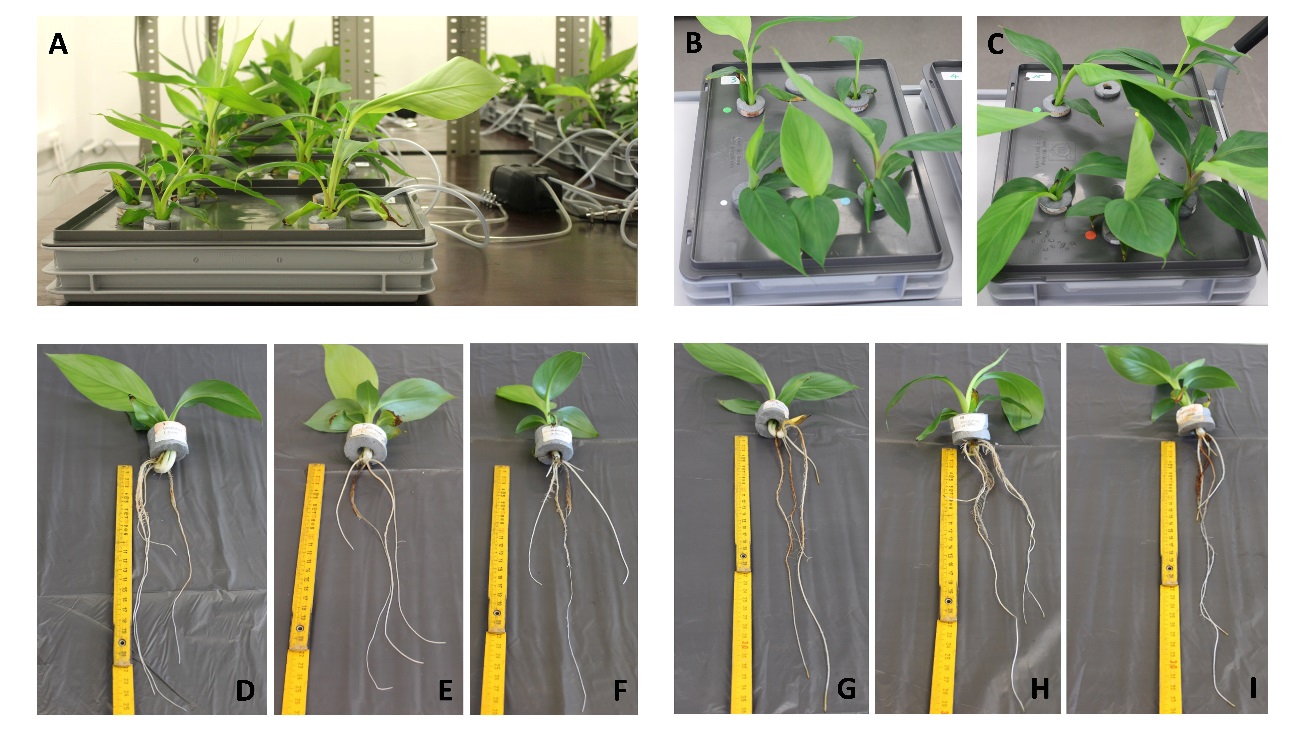

**Figure S6. Overview of the experimental set-up and plant status after 3 days of chitosan treatment in trays system. A)** Banana plants growing in trays with hydroponic medium and connected to air pumps to keep normoxic conditions ([O_2_] =7-8 mg/L). **B)** Tray with five control plants at the moment of root sampling. each plant belonging to a different banana genotype. **C)** Tray with five chitosan-treated plants at the moment of root sampling. each belonging to a different banana genotype. **D-F)** Status of control banana plants at root sampling. **G-I)** Status of chitosan-treated banana plants at root sampling.

**Supplementary tables**

**Table S1. Chitosan modifies the structure of the lipid membrane of banana plantlets roots over time.** Membrane lipids (percentage of FA with respect to the total amount of FA detected. mean ± standard error) of banana plantlets roots were analyzed after being irrigated with chitosan at different concentrations (0. 0.1 and 1 mg·ml^-1^) for 10 and 20 days (20- and 30-day-old plants). Roots of 10-day-old plants were analyzed too. to have an initial control measurement. PUFA = polyunsaturated fatty acids; n-3/n-6 = ratio between n-3 and n-6 PUFA; total lipid = % lipids in the sample.

|  | **Treatment duration (days)** | | | | | | |
| --- | --- | --- | --- | --- | --- | --- | --- |
|  | **10 day-old plants** | **10** | | | **20** | | |
|  | **Chitosan (mg·mL^-1^)** | | | | | | |
| **Fatty acids** | **[0]** | **[0]** | **[0.1]** | **[1]** | **[0]** | **[0.1]** | **[1]** |
| **14:0** | 0.18±0.03 | 0.21±0.04 | 0.15±0.05 | 0.35±0.05 | 0.14±0.07 | 0.23±0.02 | 0.26±0.04 |
| **15:0** | 0.99±0.02 | 1.14±0.07 | 1.16±0.05 | 1.31±0.07 | 1.11±0.06 | 1.21±0.06 | 1.29±0.06 |
| **16:0** | 29.06±1.31 | 29.13±1.24 | 30.23±1.21 | 29.48±0.90 | 27.83±0.36 | 28.75±0.91 | 26.47±0.79 |
| **18:0** | 6.00±0.36 | 6.13±0.36 | 5.96±0.44 | 6.16±0.25 | 5.28±0.21 | 5.96±0.35 | 5.59±0.12 |
| **20:0** | 0.99±0.06 | 1.10±0.03 | 1.10±0.10 | 1.10±0.13 | 1.15±0.08 | 1.35±0.06 | 1.16±0.15 |
| **22:0** | 0.98±0.06 | 1.02±0.10 | 1.02±0.13 | 1.12±0.19 | 1.03±0.04 | 1.31±0.16 | 0.94±0.11 |
| **24:0** | 3.79±0.10 | 4.17±0.23 | 4.29±0.33 | 3.78±0.31 | 4.60±0.16 | 4.92±0.16 | 3.27±0.47 |
| **Total saturated** | 41.98±1.85 | 42.90±1.75 | 43.91±2.06 | 43.30±1.76 | 41.14±0.52 | 43.73±1.40 | 39.00±0.82 |
| **16:1n-7** | - | 0.22±0.08 | 0.18±0.10 | 0.30±0.04 | 0.33±0.22 | 0.28±0.08 | 0.30±0.06 |
| **18:1n-9** | 2.84±0.25 | 3.04±0.09 | 2.88±0.23 | 2.44±0.14 | 3.47±0.06 | 3.06±0.25 | 2.76±0.12 |
| **18:1n-7** | 0.42±0.05 | 0.80±0.10 | 0.63±0.15 | 0.74±0.08 | 0.84±0.13 | 1.05±0.13 | 1.09±0.08 |
| **20:1n-9** | 0.21±0.01 | 0.21±0.02 | 0.21±0.01 | 0.15±0.01 | 0.21±0.01 | 0.23±0.01 | 0.15±0.00 |
| **22:1n-9** | 0.33±0.12 | 0.19±0.03 | 0.15±0.01 | 0.22±0.02 | 0.16±0.03 | 0.14±0.02 | 0.13±0.01 |
| **Total monounsaturated** | 3.81±0.32 | 4.46±0.21 | 4.05±0.39 | 3.86±0.21 | 5.01±0.26 | 4.76±0.43 | 4.43±0.02 |
| **18:2n-6** | 25.20±0.62 | 24.20±0.70 | 23.70±1.28 | 23.44±0.86 | 23.13±0.48 | 22.64±0.58 | 27.69±0.22 |
| **18:3n-6** | 0.17±0.01 | 0.16±0.01 | 0.17±0.01 | 0.17±0.00 | 0.17±0.00 | 0.16±0.00 | 0.17±0.01 |
| **20:4n-6** | 0.57±0.03 | 0.89±0.04 | 0.84±0.07 | 0.86±0.09 | 0.92±0.00 | 1.06±0.03 | 0.84±0.07 |
| **Total n-6 PUFA** | 25.94±0.64 | 25.25±0.71 | 24.71±1.29 | 24.47±0.81 | 24.22±0.48 | 23.94±0.58 | 28.70±0.15 |
| **18:3n-3** | 28.17±1.79 | 27.27±1.48 | 27.24±1.79 | 28.31±1.22 | 29.51±0.79 | 27.39±1.05 | 27.76±0.77 |
| **18:4n-3** | 0.08±0.00 | 0.08±0.01 | 0.08±0.01 | 0.06±0.00 | 0.05±0.03 | 0.09±0.01 | 0.07±0.00 |
| **20:3n-3** | - | 0.04±0.02 | 0.02±0.02 | - | 0.06±0.03 | 0.08±0.01 | 0.04±0.02 |
| **20:5n-3** | 0.03±0.03 | - | - | - | - | - | - |
| **Total n-3 PUFA** | 28.27±1.79 | 27.39±1.48 | 27.34±1.77 | 28.37±1.22 | 29.62±0.77 | 27.56±1.05 | 27.88±0.78 |
| **Total PUFA** | 54.21±2.00 | 52.64±1.91 | 52.04±2.31 | 52.84±1.83 | 53.84±0.58 | 51.51±1.57 | 56.58±0.83 |
| **n-3/n-6** | 1.09±0.07 | 1.08±0.05 | 1.12±0.09 | 1.16±0.04 | 1.22±0.05 | 1.15±0.03 | 0.97±0.03 |
| **Total lipid** | - | 0.49±0.11 | 0.71±0.18 | 0.37±0.05 | 0.53±0.15 | 0.30±0.02 | 0.47±0.06 |

**Table S2. Excitation and emission pairs of coordinates of the components for each PARAFAC model per time.** IAA = indole acetic acid. SA = salicylic acid. Component 1 includes a putative fluorophore with an Ex/Em wavelength pair of 280/368 nm. which may correspond to IAA (Li et al.. 2009). Component 2 was also induced at 5 days at the highest dose of chitosan (2 mg/mL) (Figure S1C). this component includes an Ex/Em wavelength pair of 330/416 nm. which could correspond to salicylic acid derivatives and phenolics (Street and Schenk. 1981).

| **Components**  Excitation / Emission | **1 (IAA) (Li et al.. 2009)** | **2 (SA) (Street and Schenk. 1981)** | **3 (AROMATIC AMINO ACIDS AND PEPTIDES AND PHENOLICS) (PROTEINS) (REFERENCIA)** |
| --- | --- | --- | --- |
| 1 day | 255-280 / 368 | 245 - 270 - 330 / 416 | 235 - 275 / 310 |
| 3 days | 260 - 290 / 376 | 250 - 270 - 330 / 414 | 235 - 275 / 318 |
| 5 days | 260 - 290 / 370 | 280 - 330 / 398 | - |

**Table S3. ANOVA results showing the significance level of the genotype. treatment and genotype x treatment effects for the 42 selected genes analysed by RT-qPCR.** *p < 0.05. **p < 0.01. ***p < 0.001. ****p < 0.0001. ***** p < 0.00001. ****** p < 0.000001. n.s.: not significant. Gene abbreviations according to Table S3.

| **Category** | **Gene abbreviation** | **1 day** | | | **3 days** | | |
| --- | --- | --- | --- | --- | --- | --- | --- |
|  |  | **Genotype effect** | **Treatment effect** | **Genotype × treatment effect** | **Genotype effect** | **Treatment effect** | **Genotype × treatment effect** |
|  |  | ***p*-value** | | | ***p*-value** | | |
| SA defense | *FMO1* | n.s. | ****** | * | ** | ****** | * |
|  | *EDS1* | ** | ****** | * | *** | n.s. | ** |
|  | *PAD4* | ** | ****** | * | **** | ****** | *** |
|  | *ICS1* | ** | * | n.s. | ** | ** | n.s. |
|  | *PR1* | ****** | ****** | *** | ****** | ****** | * |
|  | *NPR1* | *** | ** | * | **** | *** | * |
|  | *PAL* | ****** | ****** | * | ***** | n.s. | * |
|  | *PPO* | * | ****** | n.s. | ** | ***** | n.s. |
| Auxins | *AAO1* | n.s. | ** | n.s. | * | * | n.s. |
|  | *AMI1* | ***** | *** | * | ****** | * | n.s. |
|  | *ARF1* | * | n.s. | n.s. | n.s. | ****** | n.s. |
|  | *TAA1* | ** | n.s. | n.s. | *** | ** | ** |
|  | *PIN1* | ***** | ** | n.s. | **** | * | n.s. |
|  | *YUCCA2* | ***** | ****** | n.s. | ****** | ****** | * |
| Hypoxia | *AOX* | ***** | ****** | ** | *** | ****** | n.s. |
|  | *HB* | n.s. | ****** | n.s. | ** | *** | n.s. |
|  | *ADH* | ** | ***** | n.s. | *** | n.s. | n.s. |
|  | *LDH* | *** | ****** | ** | n.s. | n.s. | n.s. |
|  | *PDC* | n.s. | ****** | n.s. | n.s. | n.s. | n.s. |
| ROS detoxification | *APX* | ****** | n.s. | n.s. | ****** | * | n.s. |
|  | *CAT* | ****** | * | *** | ****** | n.s. | n.s. |
|  | *ERO1* | ** | ** | n.s. | *** | ** | n.s. |
|  | *GR* | **** | * | n.s. | ****** | ** | n.s. |
|  | *SOD* | **** | ** | n.s. | ****** | *** | n.s. |
| Ethylene & ABA | *ACO* | *** | ***** | n.s. | * | ***** | n.s. |
|  | *ERF1* | ****** | * | * | ****** | * | n.s. |
|  | *ASA1* | ****** | n.s. | n.s. | ****** | * | n.s. |
|  | *NCED3* | * | ** | n.s. | n.s. | ** | n.s. |
| JA defense | *AOS* | **** | **** | n.s. | n.s. | * | n.s. |
|  | *AOC* | ****** | ****** | *** | ***** | * | n.s. |
|  | *COI1* | *** | ** | n.s. | ** | **** | * |
|  | *JMT* | ****** | ***** | n.s. | ***** | n.s. | * |
|  | *MYC2* | **** | *** | * | **** | *** | n.s. |
| Root growth & development | *AQC1* | ** | n.s. | n.s. | ** | **** | n.s. |
|  | *LOX* | ** | ****** | n.s. | * | n.s. | n.s. |
|  | *WOX5* | *** | **** | n.s. | *** | n.s. | ** |
|  | *GLP10* | n.s. | n.s. | n.s. | n.s. | ***** | * |
| Parasitism & endophytism | *APP* | *** | n.s. | n.s. | **** | ****** | n.s. |
|  | *AP1* | n.s. | * | n.s. | **** | ** | * |
|  | *CytP450* | *** | * | n.s. | ****** | n.s. | n.s. |
|  | *NRT1* | ****** | * | n.s. | ***** | n.s. | n.s. |
